# An orthologous gene coevolution network provides insight into eukaryotic cellular and genomic structure and function

**DOI:** 10.1101/2021.07.09.451830

**Authors:** Jacob L. Steenwyk, Megan A. Phillips, Feng Yang, Swapneeta S. Date, Todd R. Graham, Judith Berman, Chris Todd Hittinger, Antonis Rokas

**Affiliations:** Vanderbilt University, Department of Biological Sciences, Nashville, TN, United States of America; Shmunis School of Biomedical and Cancer Research, Tel Aviv University, Ramat Aviv, Israel; Department of Pharmacology, Shanghai Tenth People’s Hospital, Tongji University School of Medicine, Shanghai, China; Laboratory of Genetics, DOE Great Lakes Bioenergy Research Center, Wisconsin Energy Institute, Center for Genomic Science Innovation, J.F. Crow Institute for the Study of Evolution, University of Wisconsin-Madison, Madison, Wisconsin, United States of America

**Keywords:** Evolutionary rate covariation, DNA replication, DNA repair, genetic network, chromatin remodeler, Golgi transport

## Abstract

Orthologous gene coevolution—which refers to gene pairs whose evolutionary rates covary across speciation events—is often observed among functionally related genes. We present a comprehensive gene coevolution network inferred from the examination of nearly three million orthologous gene pairs from 332 budding yeast species spanning ∼400 million years of eukaryotic evolution. Modules within the network provide insight into cellular and genomic structure and function, such as genes functioning in distinct cellular compartments and DNA replication. Examination of the phenotypic impact of network perturbation across 14 environmental conditions using deletion mutant data from the baker’s yeast *Saccharomyces cerevisiae* suggests that fitness in diverse environments is impacted by orthologous gene neighborhood and connectivity. By mapping the network onto the chromosomes of *S. cerevisiae* and the opportunistic human pathogen *Candida albicans*, which diverged ∼235 million years ago, we discovered that coevolving orthologous genes are not clustered in either species; rather, they are most often located on different chromosomes or far apart on the same chromosome. The budding yeast coevolution network captures the hierarchy of eukaryotic cellular structure and function, provides a roadmap for genotype-to-phenotype discovery, and portrays the genome as an extensively linked ensemble of genes.

## Introduction

Genetic networks—diagrams wherein nodes represent genes and edges represent measured functional relationships between nodes—can elucidate how genes are organized into pathways and contribute to cellular functions, shedding light into the relationship between genotype and phenotype (Costanzo *et al*. 2010, 2016, 2019; Kuzmin *et al*. 2018). Given the rich information contained in or derived from genetic networks, numerous approaches that aim to capture some aspect(s) of functional relationships among genes in a genome (e.g., gene coexpression, genetic interaction) have been developed (Lezon *et al*. 2006; Baryshnikova *et al*. 2013; Wisecaver *et al*. 2017). While these networks are highly informative, their availability and applicability is typically limited to select model organisms and single extant species or strains. Application of information from the genetic network of one organism to understand the biology of another requires assuming that the networks of the two organisms are conserved, which is not always the case (Tong *et al*. 2001; Pan *et al*. 2004; Onge *et al*. 2007; Mani *et al*. 2008; Dixon *et al*. 2008; Lehner 2011; Boucher and Jenna 2013; Lind *et al*. 2015; Sorrells and Johnson 2015; Monaco *et al*. 2015; Yang and Wittkopp 2017).

One complementary, but poorly studied, method for constructing genetic networks is by measuring the coevolution of orthologous genes, which can be done by calculating the covariation of relative evolutionary rates among orthologous genes (Goh *et al*. 2000; Sato *et al*. 2005; Clark *et al*. 2012; Steenwyk *et al*. 2021). Briefly, by estimating an orthologous gene’s phylogeny, one infers the rate (and changes in rate) of its evolution across the phylogeny; if the evolutionary rate values estimated for each branch of an orthologous gene’s phylogeny are significantly correlated with those of another gene’s phylogeny, the two orthologs are said to be coevolving. By estimating coevolution for all pairs of orthologous genes in a clade, one can infer the clade’s orthologous gene coevolution network, where nodes correspond to orthologs and edges correspond to the degree to which two orthologs coevolve (Steenwyk *et al*. 2021). Genetic networks based on gene coevolution leverage evolutionary information, whereas standard genetic networks rely on the correlation of functional data such as gene expression or the presence of genetic interactions among genes within a single extant species or strain.

Orthologous gene coevolution is often observed among genes that share functions, are coexpressed, or whose protein products are subunits in a multimeric protein structure, and can yield insights into the genotype-to-phenotype map (Findlay *et al*. 2014; Brunette *et al*. 2019). For example, screening for genes that have coevolved with genes in known DNA repair pathways across 33 mammals led to the identification of *DDIAS*, whose involvement in DNA repair was subsequently functionally validated (Brunette *et al*. 2019). Furthermore, four out of five proteins in the protein structural interactome map—a database of structural domain-domain interactions in the protein data bank (https://www.rcsb.org/)—exhibit signatures of gene coevolution (Kim *et al*. 2004). Although these and other studies have demonstrated that signatures of coevolution are a powerful method to detect functional associations among genes in the absence of functional data (Clark *et al*. 2012; Findlay *et al*. 2014; Raza *et al*. 2019; Brunette *et al*. 2019; Huang *et al*. 2020; Talsness *et al*. 2020), the network biology principles of gene coevolution, especially between genes that have coevolved for hundreds of millions of years, remain unexplored.

To unravel general principles of orthologous gene coevolutionary networks, we constructed the coevolution network of a densely sampled set of orthologs from one-third of known budding yeast species (332 species) that diversified over ∼400 million years. The inferred network provides a hierarchical view of cellular function from broad bioprocesses to specific pathways. Interpolation of the gene coevolution network with of fitness assay data from single- and digenic *S. cerevisiae* mutants (Costanzo *et al*. 2010, 2016, 2021; Usaj *et al*. 2017) provides insight into subnetwork- and ortholog-specific potential to buffer genetic perturbations. Surprisingly, comparisons of genetic networks inferred from gene coevolution and genetic interactions yield similar functional insights; for example, hubs of genes tend to be functionally related and gene essentiality impacts gene connectivity wherein essential genes are more densely connected than non-essential genes. Unlike genetic interaction networks, gene coevolution networks can also provide evolutionary insights; for example, mapping the orthologous gene coevolution network onto the chromosomes of two model yeast genomes uncovers extensive inter-chromosomal and long-range intra-chromosomal associations, providing an ‘entangled’ view of the genome across evolutionary timescales. We anticipate these results will facilitate the generation, interpretation, and utility of these networks among other lineages in the tree of life.

## Results

### A gene coevolution network

We examined 2,898,028 pairs of orthologous genes from a dataset of 2,408 orthologous gene in 332 budding yeast species. Broad network properties were stable across a range of thresholds for “significant” orthologous gene coevolution (Fig. S2). To conservatively define “significant” coevolution and therefore examine orthologous gene pairs with only robust signatures of coevolution, we implemented a high correlation coefficient threshold for significant orthologous gene coevolution (r ≥ 0.825; Pearson correlation among relative evolutionary rates). This resulted in 60,305 significant signatures of orthologous gene coevolution; Fig 1A, 1B, and S1), which were used to construct a network where nodes are orthologous genes and edges connect orthologous genes that are significantly coevolving (Fig. 1C).

**Figure 1.**
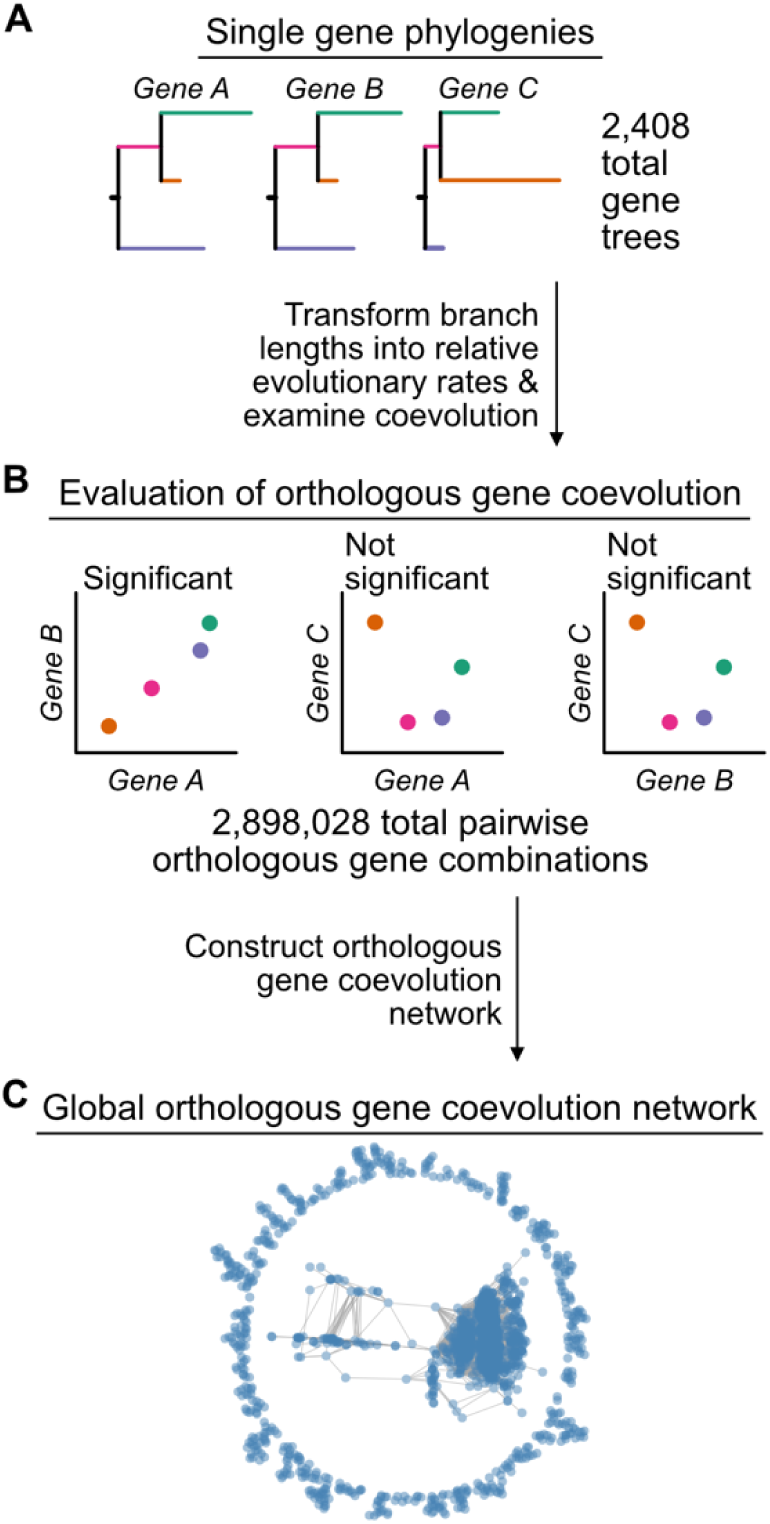
Constructing the budding yeast orthologous gene coevolution network. (A) We determined coevolution in a set of 2,408 single gene trees in which branch lengths were inferred along the species tree topology. (B) Coevolution of orthologous genes was evaluated across all pairwise combinations of orthologous genes using the CovER function in PhyKIT, v0.1 (Steenwyk *et al*. 2021). (C) Significantly coevolving pairs of orthologous genes were used to construct a global network of orthologous gene coevolution where nodes correspond to orthologous genes and edges connect orthologous genes that are significantly coevolving.

To determine how orthologous gene connectivity varied in the network, we examined patterns of dense and sparse connections for individual orthologous genes. Individual orthologous genes coevolved with a median of eight other orthologous genes, but connectivity varied substantially across the network (Fig. S3). For example, 1,091 orthologous genes have signatures of coevolution with five or fewer other orthologous genes and 601 orthologous genes are singletons (i.e., they are not significantly coevolving with other orthologous genes in the dataset). In contrast, 420 orthologous genes have signatures of coevolution with 100 or more other orthologous genes, and 21 orthologous genes coevolve with 400 or more others.

Coevolving orthologous genes in the network are functionally related. For example, *PEX1* and *PEX6* are one of the pairs of genes with the highest observed correlation coefficient in evolutionary rates (Fig. S4). In *S. cerevisiae*, the two orthologous genes encode a heterohexameric complex responsible for protein transport across peroxisomal membranes (Ciniawsky *et al*. 2015) and mutations in either gene can lead to severe peroxisomal disorders in humans (Reuber *et al*. 1997). Functional enrichment among densely connected orthologous genes revealed that complex bioprocesses that require coordination among polygenic protein products are overrepresented (Fig. S5, Table S1). For example, *CHD1, INO80*, and *ARP5*, which encode proteins responsible for chromatin remodelling processes such as nucleosome sliding and spacing (Ayala *et al*. 2018), are coevolving with 400 or more other orthologous genes (Fig. S5, Table S1). Taken together, these findings highlight that coevolution may be observed among orthologous genes that physically interact (e.g., *PEX1* and *PEX6*) or contribute to highly intricate biological processes (e.g., *INO80*). More broadly, these data support the hypothesis that coevolving orthologous genes tend to have similar functions.

To determine how connectivity varied within the network, we examined the properties of subnetworks across orthologous genes considered essential and nonessential in the model yeast *S. cerevisiae* or the opportunistic pathogen *C. albicans* (Winzeler *et al*. 1999; Segal *et al*. 2018). Essential genes are densely connected in the orthologous gene coevolutionary network, whereas nonessential genes exhibit sparser connections (Fig. 2A-D). To infer network communities— clusters of orthologous genes that have more connections between them than between orthologous genes of different clusters—we used a hierarchical agglomeration algorithm (Fig. 2A). Five large communities (clusters of more than 10 orthologous genes) were identified. Each community varied in size, community-to-community connectivity, and essential/nonessential orthologous gene composition. Specifically, the two largest communities, communities 1 and 2, share the most connections and belong to a higher-order cluster with the next two largest communities, communities 3 and 4 (Fig. 2E and S6). In contrast, the smallest community, community 5, does not cluster with the other communities. Similarly, essential genes are overrepresented in community 1 but are underrepresented in communities 2, 3, and in smaller communities of 10 or fewer orthologous genes (Fig. 2F; p < 0.01 for all tests; Fisher’s exact test). The result that *S. cerevisiae* and *C. albicans* essential genes are central hubs in coevolution network constructed from orthologous genes that represent 400 million years of budding yeast evolution mirrors observations from the *S. cerevisiae* genetic interaction network (Costanzo *et al*. 2016).

**Figure 2.**
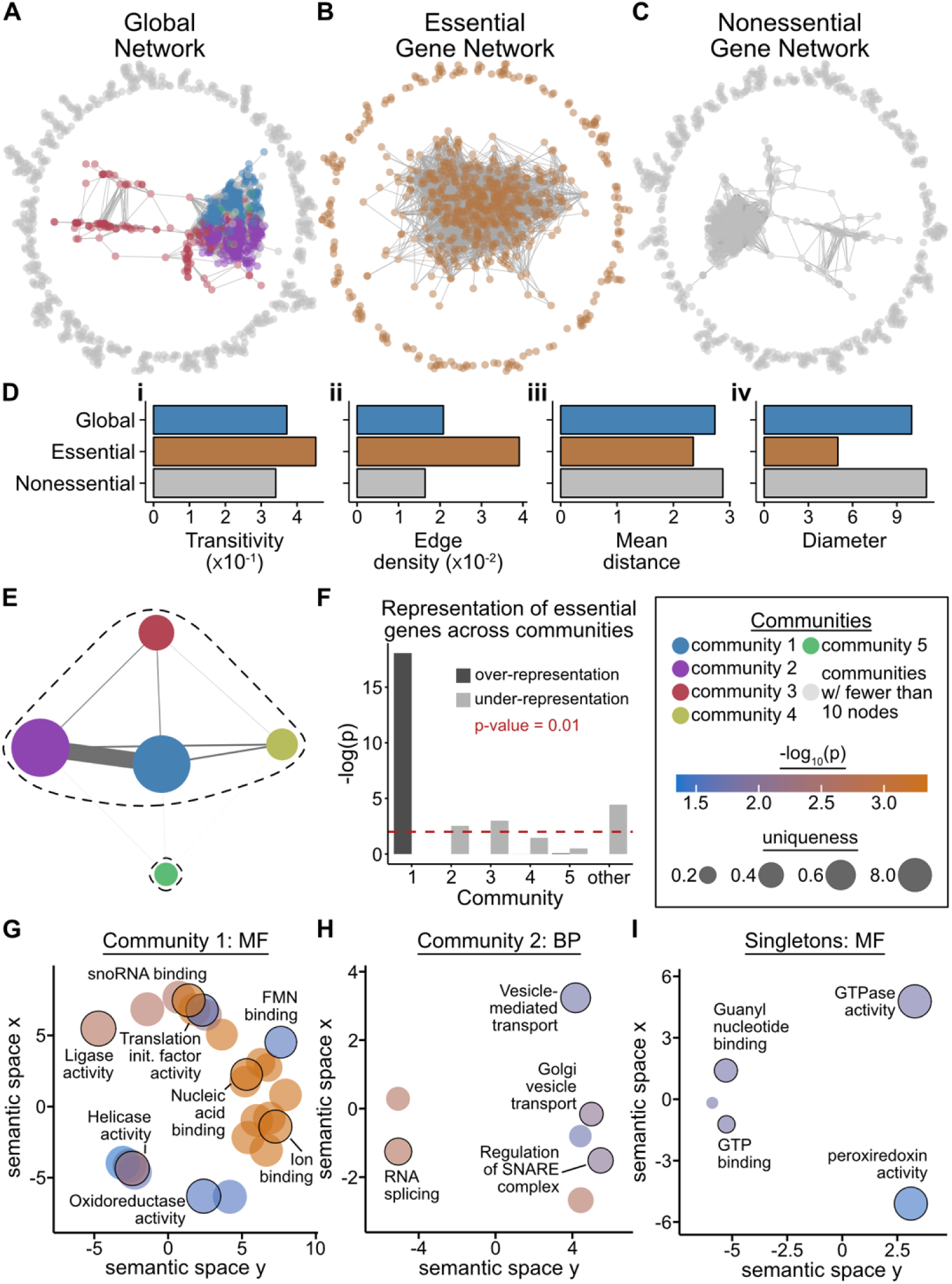
Network modules reflect modules of bioprocesses. (A) The global network of orthologous gene coevolution. Subnetworks of orthologous genes that are essential (B) and nonessential (C) in *S. cerevisiae* and *C. albicans*. (D) Examination of network properties reveals that the essential gene subnetwork has higher values for metrics of network density (i.e., transitivity and edge density), whereas the nonessential gene network has higher values for metrics that measure how diffuse the network is (i.e., mean distance and diameter). (E) Network community detection revealed five major subnetworks or communities. Orthologous genes from each community are depicted in the same color in panel A and genes from small communities with 10 or fewer orthologous genes are depicted in gray. There are 804, 740, 161, 39, and 15 genes in the five largest communities (communities 1-5). Edge width reflects the number of co-evolving orthologous gene pairs between the two communities and node size reflects the number of orthologous genes in each community. Higher-order community clustering revealed communities 1 through 4 cluster together, whereas community 5 is a singleton as denoted by the dashed line. (F) Community 1 is overrepresented with orthologs whose genes are essential in *S. cerevisiae* and *C. albicans* whereas all other communities are underrepresented with essential genes. (G-I) Communities differ in enriched terms; for example, enriched molecular functions in community 1 includes orthologous genes associated with helicase, ligase, and translation initiation factor activities. MF and BP represent molecular functions and biological processes, respectively. Each enriched GO term is represented by a circle where circle color reflects -log_10_ p-value from a Fischer’s exact test with Benjamini Hochberg multi-test correction and the size of each circle represents GO term uniqueness, a measure of GO term dissimilarity to other enriched GO terms wherein higher values reflect greater uniqueness. Complete enrichment results for each community are reported in Table S3. The box to the right of panel F is the legend for the whole figure.

### From processes to pathways: the budding yeast coevolution network captures the hierarchy of cellular function

To gain insight into the functional neighborhoods of the orthologous gene coevolution network, we examined via gene ontology (GO) enrichment analysis (GeneOntologyConsortium 2004) the composition of each community. Among the highest-order cluster of communities (i.e., communities 1 through 4), we found that higher-order cellular processes including nucleic acid metabolism (p = 0.040; Fisher’s exact test multi-test corrected using false discovery rate correction with Benjamini/Hochberg (FDR-BH)) and cellular anatomical entities (p = 0.020; Fisher’s exact test multi-test corrected using FDR-BH) are enriched. At the individual community level, we found that community 1 is enriched in orthologous genes with helicase activity (p = 0.005; Fisher’s exact test multi-test corrected using FDR-BH), ligase activity (p = 0.004; Fisher’s exact test multi-test corrected using FDR-BH), and translation initiation factors (p = 0.024; Fisher’s exact test multi-test corrected using FDR-BH); community 2 is enriched in Golgi vesicle transport orthologous genes (p = 0.009; Fisher’s exact test multi-test corrected using FDR-BH); whereas singletons are enriched in GTPase activity (p = 0.016; Fisher’s exact test multi-test corrected using FDR-BH) and peroxiredoxin activity (p = 0.036; Fisher’s exact test multi-test corrected using FDR-BH) (Fig. 2G-I, Table S3).

Functional neighborhoods of coevolving orthologous genes within and between biological functions as well as cellular compartments and complex categories are also captured by the network. For example, orthologous genes involved in the biological functions of ribosome biogenesis, rRNA processing, and translation, which represent different functional categories, are extensively coevolving with one another (Fig. S7A). This finding suggests that the complexity of protein biosynthesis, a process that requires coordination among diverse biochemical functions, is captured in the coevolution of the underlying orthologous genes. Similarly, orthologous genes involved in nuclear processes or located in the cytoplasm tend to coevolve with orthologous genes in the same cellular compartment, however, substantial signatures of coevolution between orthologous genes from different cellular compartments are also observed (Fig. S7B).

Finally, our network captures functional neighborhoods of coevolving orthologous genes at the level of pathways and complexes. We found strong signatures of coevolution among orthologous genes from specific pathways and complexes. For example, orthologous genes that encode proteins responsible for DNA replication coevolve with a larger number of other DNA replication orthologous genes than expected by random chance (p < 0.001; permutation test) (Fig. S8). Orthologous genes involved in DNA mismatch repair and nucleotide excision repair pathways, which participate in the repair of DNA lesions, have more signatures of coevolution than expected by random chance (p < 0.001 for each pathway; permutation test). Orthologous genes in the phosphatidylcholine biosynthesis pathway, which is responsible for the biosynthesis of the major phospholipid in organelle membranes, and orthologous genes in the tricarboxylic acid cycle (also known as the Krebs cycle or citric acid cycle), a key component of aerobic respiration (Fig. S9), also have more signatures of coevolution than expected by random chance (p < 0.001 for each pathway; permutation test). Among complexes, orthologous genes that encode the minichromosome maintenance protein complex that functions as a DNA helicase, the DNA polymerase α-primase complex that assembles RNA-DNA primers required for replication, and DNA polymerase ε that serves as a leading strand DNA polymerase (Fig. 3) also coevolve with larger numbers of orthologs from the same complex than expected by random chance (p < 0.001 for each multimeric complex; permutation test). Note, certain gene categories (e.g., transposons and hexose transporters) are not represented in our dataset of orthologous genes and could not be examined (see *Methods*).

**Figure 3.**
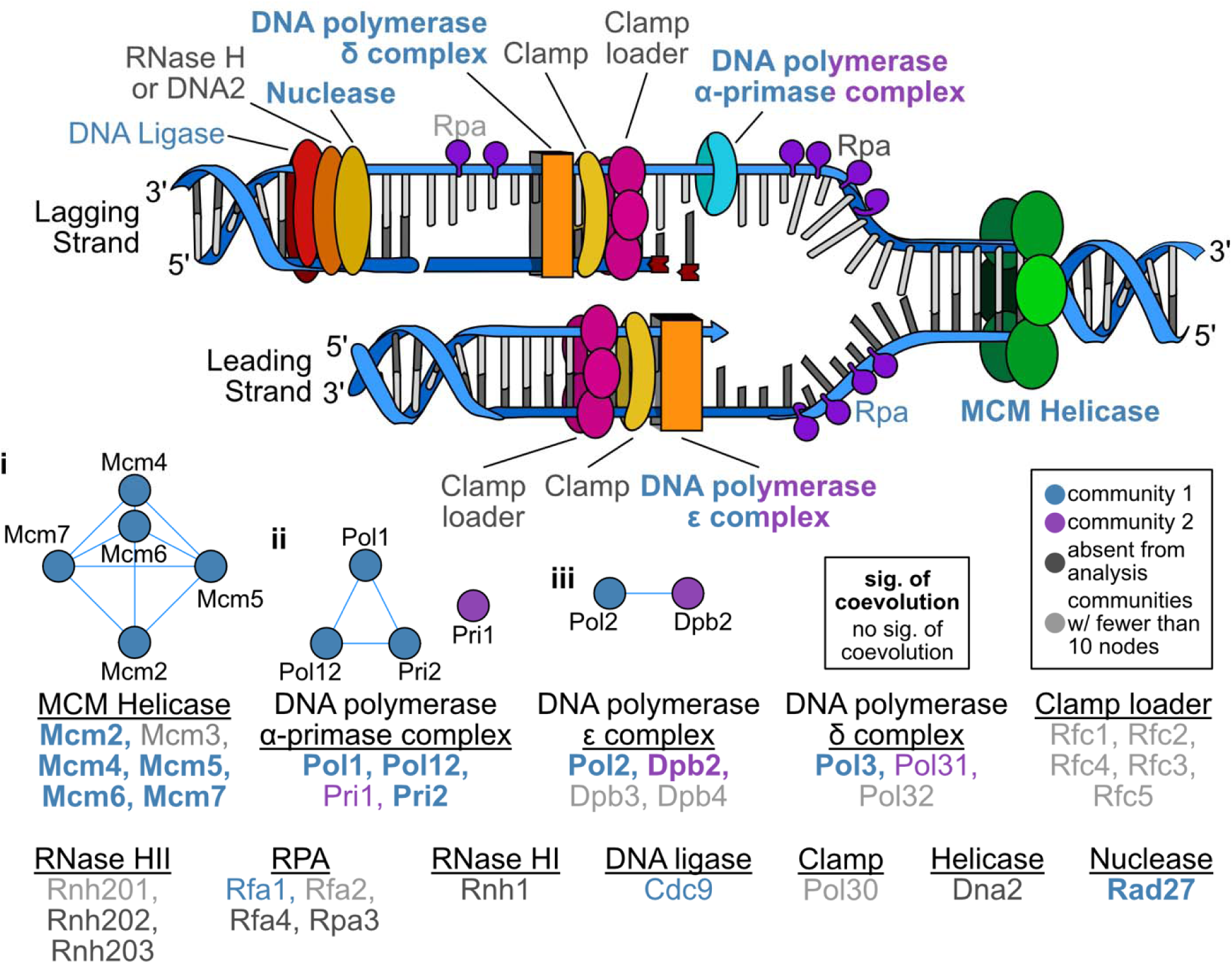
Extensive coevolution in DNA replication genes. Cartoon representation of DNA replication. Exemplary complex specific subnetworks are depicted in i, ii, and iii. (i) Extensive coevolution between orthologous genes that encode the helicase, minichromosome maintenance (MCM) complex, which functions as a helicase. (ii) Coevolution in the orthologous genes that encode the DNA polymerase α-primase complex and (iii) DNA polymerase ε complex, which are responsible for RNA primer synthesis and leading strand DNA synthesis, respectively. Edges in blue connect orthologous genes that are significantly coevolving. Orthologous genes and complexes in bold have signatures of coevolution. Orthologous genes and complexes are colored according to community assignment. Complexes, such as the DNA polymerase α-primase complex, are depicted in multiple colors reflecting the multiple communities represented within the complex. There is significant coevolution across all DNA replication orthologous genes (p < 0.001; permutation test) as well as the multimeric complexes such as the MCM complex (p < 0.001 for each pathway; permutation test).

In summary, these findings reveal that functional aspects of the network can be viewed with varying degrees of specificity. For example, the highest-order insights (i.e., GO enrichment across communities 1, 2, 3, and 4) revealed coevolution among cellular anatomical entities whereas greater specificity—such as coevolution among orthologous genes responsible for Golgi vesicle transport—can be obtained by examining lower-order hubs of genes (e.g., GO enrichment in community 2). Furthermore, coevolutionary signatures can bridge distinct but related functional categories such as cellular compartments and complexes, highlighting the complex interplay of distinct functional modules over evolutionary time. Thus, the budding yeast coevolution network captures the hierarchy of cellular function from broad bioprocesses to specific pathways or multimeric complexes.

### The coevolution network constructed from budding yeast orthologous genes is distinct, but complementary, to the *S. cerevisiae* genetic interaction network

To determine similarities and differences between our coevolution network inferred from orthologous genes in the budding yeast subphylum and the genetic interaction network inferred from digenic null mutants in the model organism *S. cerevisiae* (Costanzo *et al*. 2010; Usaj *et al*. 2017), both data types were integrated into a single supernetwork (Fig. S10 and S11). We hypothesize that there will be broad similarities between the networks because they both capture functional associations; however, we also hypothesize that the connectivity of individual nodes between the networks will sometimes differ because one network is built from ∼400 million years of orthologous gene coevolution whereas the other from genetic interactions in a single extant species.

Supporting this hypothesis, the community clustering observed in the gene coevolution network was also evident in the supernetwork; however, gene- / ortholog-wise connectivity at times differed suggesting each network harbors distinct and complementary insights (Fig. S10). For example, connectivity is similar for the gene / ortholog *CDC6*, which is required for DNA replication (Hartwell *et al*. 1970), between the two networks. Specifically, *CDC6* is connected to 96 genes / orthologs in both networks and 56 of the genes / orthologs are the same. This result suggests that the connectivity of the *CDC6* gene in *S. cerevisiae* is broadly conserved across species from the budding yeast subphylum. In contrast, different gene- / ortholog-wise connectivity was observed for the choline kinase *CKI1* (Hosaka *et al*. 1989; Kim *et al*. 1999); *CKI1* is coevolving with 87 orthologs, has a significant genetic interaction with 10 genes, and seven of these genes / orthologs are shared by both networks. This result suggests that the connectivity of the *CKI1* gene observed in *S. cerevisiae* is not broadly conserved across species from the budding yeast subphylum. This difference may be partially explained by the fact that *CKI1* has a paralog, *EKI1*, which arose from an ancient whole genome duplication event that affected some, but not all, species in the subphylum (Marcet-Houben and Gabaldón 2015; Wolfe 2015). These results reveal that orthologous gene coevolution networks inferred over macroevolutionary timescales and networks inferred from genetic interactions in single organisms offer complementary insights into functional relationships between genes.

### Communities differ in capacity to compensate for perturbation

Examinations of genome-wide gene dispensability in the model budding yeast *S. cerevisiae* and the opportunistic pathogen *Candida albicans* (Winzeler *et al*. 1999; Segal *et al*. 2018) suggest that single-organism genetic networks can buffer perturbations resulting from the deletion of individual genes. Thus, we sought to determine whether a gene’s dispensability varies in a community-dependent manner. To address this, we integrated information from the budding yeast orthologous gene coevolution network and genome-wide single-gene deletion fitness assays (or, in the case of essential genes, expression suppression) of *S. cerevisiae* in 14 diverse environments (Costanzo *et al*. 2021) (Fig. S12 and S13). Here, single-gene deletion fitness assays serve as a proxy for network perturbation in which deletion of a single gene is analogous to removing a node from the network. We found that fitness of *S. cerevisiae* gene knockouts in different environments was significantly dependent on community and the number of coevolving genes per gene (Fig. 4; p < 0.001 for both comparisons, Multi-factor ANOVA). These observations support previous findings that the impact of single-gene deletions can be buffered but also highlight the importance of the architecture of the underlying genetic network.

**Figure 4.**
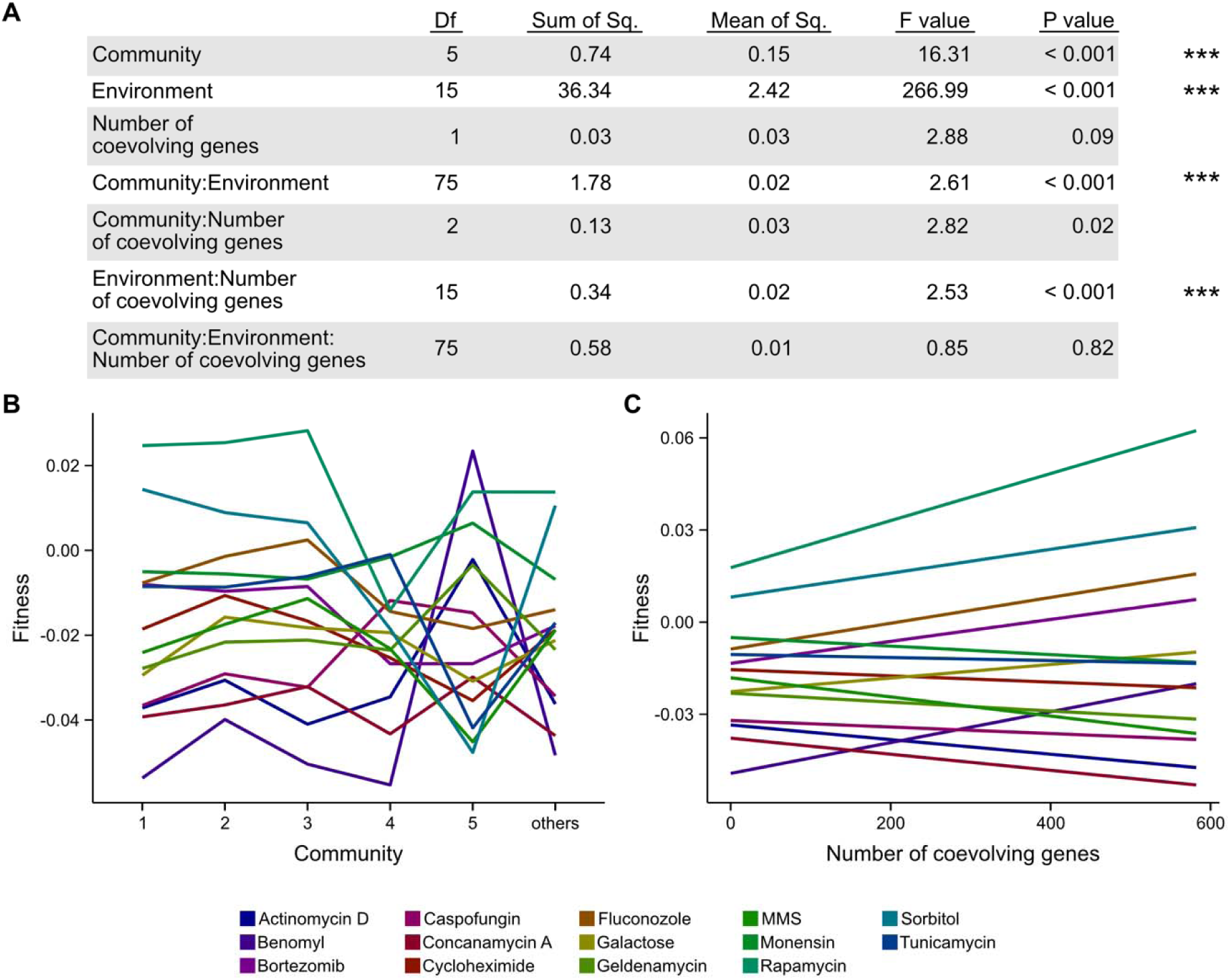
The impact of perturbing the orthologous gene coevolutionary network through single-gene deletion in diverse environments is dependent on community and gene connectivity. (A) Multi-factor ANOVA results indicate community, environment, the interaction between community and environment, and the interaction between environment and the number of coevolving orthologous genes per orthologous gene are significantly associated with the fitness of a single-gene deletion strain (relative to the wild-type strain). (B) Fitness of single-gene deletion strains in diverse environments is impacted by community. (C) Similarly, fitness of single-gene deletion strains in diverse environments is impacted by the number of coevolving orthologous genes the deleted node is connected to. These results indicate that fitness in diverse environments is impacted by orthologous gene neighborhood and connectivity in the network. In both panels, each color corresponds to a different environment that fitness was measured in. Df represents degrees of freedom; Sum of Sq. represents sum of squares; Mean of Sq. represents Mean of squares.

To further investigate the relationship between *S. cerevisiae* gene dispensability and structure of the coevolution network, we integrated *S. cerevisiae* genetic interaction data from double-gene or digenic deletion fitness assays, wherein positive and negative genetic interactions refer to positive and negative fitness effects in the digenic deletion mutants relative to those expected from the combined effects of the individual single-gene deletion mutants, respectively (Costanzo *et al*. 2010, 2016; Usaj *et al*. 2017). Although most digenic deletions were associated with negative genetic interactions, we unexpectedly found more instances of positive fitness effects for orthologous genes from the same community (e.g., deleting two genes whose orthologous genes are both in community 1) than for orthologous genes from different communities; the sole exception was digenic gene losses of orthologous genes in communities 1 and 2, which are highly connected (Fig. S14A). These results suggest that losses of two orthologous genes from the same community are difficult to buffer, but, if both orthologous genes are in the same subnetwork (or community), compensatory effects are more likely to be observed.

Finally, to examine evolutionary gene loss in the context of the gene coevolution network, we investigated community-wide patterns of gene losses among genes lost in a lineage of budding yeasts previously reported to have undergone extensive gene losses (Steenwyk *et al*. 2019). These analyses revealed community 2 and singleton orthologs are more likely to be lost (Fig. S14B), which supports the hypothesis that gene losses do not occur stochastically (Albalat and Cañestro 2016). In summary, the architecture of the coevolution network is significantly associated with a gene’s dispensability.

### An entangled genome: extensive inter- and long-range intra-chromosomal coevolution

Gene order is not random among eukaryotes and physically linked genes tend to be involved in the same metabolic pathway or protein-protein complex (Hurst *et al*. 2004; Rokas *et al*. 2018). Thus, we hypothesized that coevolving orthologous genes will likely be physically linked or clustered onto yeast chromosomes. To test this hypothesis, we projected the budding yeast gene coevolution network onto the one-dimensional genome structure of *S. cerevisiae* and *C. albicans*, which diverged ∼235 million years ago (Shen *et al*. 2018). We chose the genomes of these two organisms because they both have complete and high-quality chromosome-level assemblies. The two organisms also have distinct evolutionary histories; the lineage that includes *S. cerevisiae* underwent whole-genome duplication, whereas *C. albicans* underwent intra-species hybridization (Marcet-Houben and Gabaldón 2015; Mixão and Gabaldón 2020). These processes have contributed to differences in chromosome number (16 in *S. cerevisiae* vs. eight in *C. albicans*) and a lack of macrosynteny (Seoighe *et al*. 2000; Chibana *et al*. 2005; Wolfe 2006; Fitzpatrick *et al*. 2010; Dujon 2010) (Fig. 5A-B and Fig. S15-S16).

**Figure 5.**
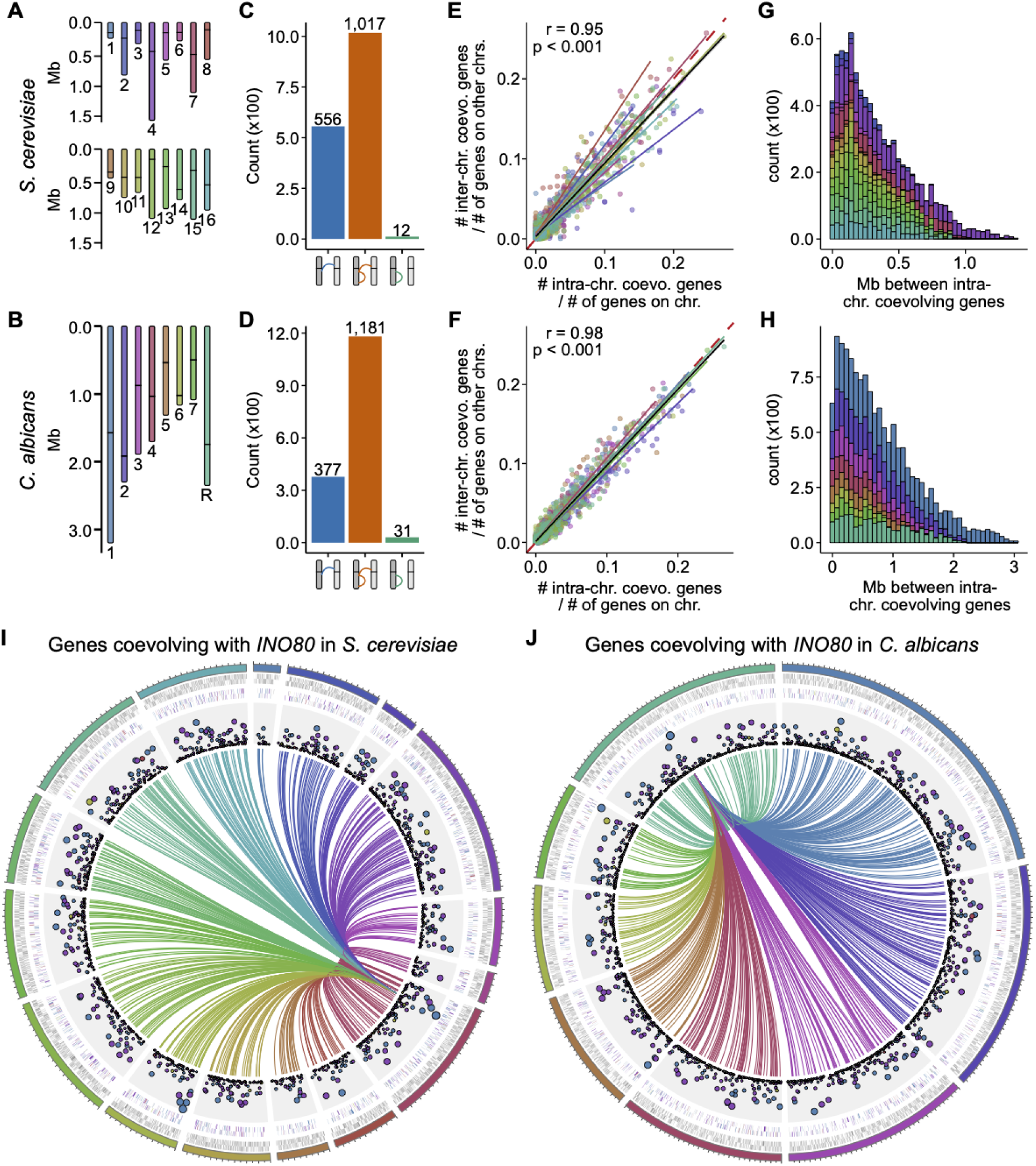
Extensive long range and inter-chromosomal gene coevolution. (A & B) The number and size of chromosomes differ in (A) *S. cerevisiae* and (B) *C. albicans*. Black lines indicate locations of centromeres. (C & D) The numbers of genes with only inter-chromosomal (blue), both inter-and intra-chromosomal (orange), or only intra-chromosomal (green) signatures of orthologous gene coevolution reveal substantially more inter-chromosomal associations than intra-chromosomal ones. (E & F) The relationship between the number of intra-chromosomal signatures of orthologous gene coevolution corrected by the number of genes on the same chromosome (x-axis) and the number of inter-chromosomal signatures of orthologous gene coevolution corrected by the number of genes on other chromosomes (y-axis) reveals intra- and inter-chromosomal associations are equally likely, suggesting function, rather than genetic neighborhood, is the primary driver of orthologous gene coevolution. Each color represents a different chromosome. Regression lines are depicted for each chromosome using a linear model. A summary regression line is depicted in black. (G & H) Distribution of distances among intra-chromosomal signatures of orthologous gene coevolution reveals long-range coevolution is common. (I & J) An exemplary orthologous gene, *INO80*, reveals how a single orthologous gene can be coevolving with other orthologous genes on the same or other chromosomes. The first, innermost track depicts the various chromosomes of either yeast in which chromosome 1 is shown at the 12 o’clock position and increasing chromosome number is depicted in a clock-wise manner. The second track shows genes on the plus or minus strand. The third track shows the same data for the genes present in our dataset and each gene is colored according to the community they are part of. The scatter plot shows the number of coevolving orthologous genes per orthologous gene. Larger circles represent orthologous genes that are connected to more orthologous genes in the network. The links depict orthologous genes that are coevolving with *INO80* and each link is colored according to the chromosomal location of the other orthologous gene it is coevolving with. Colors in E-H as well as ideogram and link colors in J correspond chromosomes as depicted in A and B.

Contrary to our hypothesis, we observed extensive inter-chromosomal and long-range intra-chromosomal orthologous gene coevolution (Fig. 5 and Fig. S17-S23). Specifically, co-evolving orthologous gene pairs were commonly located on different chromosomes (Fig. 5C-D and Table S4). There was a near-perfect correlation between the number of intra-chromosomal signatures of coevolution (corrected by the number of genes on that chromosome in the dataset) and the number of inter-chromosomal signatures of coevolution (corrected by the number of genes on all other chromosomes in the dataset) (r = 0.95, p < 0.001 for *S. cerevisiae*; r = 0.98, p < 0.001 for *C. albicans*; Spearman correlation). This result suggests that orthologous genes located on the same or different chromosomes are equally like to be coevolving. Given the extensive coevolution among orthologous genes in the same or similar functional categories, these results support the notion that function, not chromosome structure, is the primary driver of coevolution over macroevolutionary timescales.

Examination of intra-chromosomal coevolution revealed variation in orthologous gene pair distances along the genome. Two coevolving orthologous genes on the same chromosome can be kilobase-to-megabase distances from one another (Fig. 5G-H). The distribution of the closest distance between an orthologous gene and its coevolving partners revealed a positively skewed distribution with a similar range of kilobase-to-megabase associations (Fig. S23). In *S. cerevisiae*, the number of intra-chromosomal signatures of coevolution is correlated with the number of genes on a chromosome represented in the dataset, whereas in *C. albicans* the number of intra-chromosomal signatures of coevolution is correlated both with chromosome length and with the number of genes on a chromosome represented in the dataset (Fig. S24). Examination of the distances between orthologous genes in our dataset and their coevolving partners revealed that long-range intra-chromosomal coevolution was not an artifact of gene sampling (Fig. S24). Investigation of the interplay between orthologous gene coevolution and chromosomal contacts using a three-dimensional model of the *S. cerevisiae* genome (Duan *et al*. 2010) revealed signatures of coevolution occur independent of chromosomal contacts (Fig. S26).

Extensive inter- and intra-chromosomal associations are exemplified by *INO80*, which encodes a chromatin remodeler and has coevolved with 591 orthologous genes on all other chromosomes in both *S. cerevisiae* and *C. albicans* (Fig. 5I-J). To date, few examples of inter-chromosomal associations between loci are known. One example includes concerted copy number variation between 45S and 5S rDNA loci in humans; imbalance in copy number is thought to be associated with disease (Gibbons *et al*. 2014, 2015). Our observations suggest extensive inter-chromosomal and long-range intra-chromosomal functional associations may be more common than previously appreciated.

## Discussion

We constructed a genetic network based on orthologous gene coevolution from a densely sampled set of orthologs across the budding yeast subphylum. These analyses are distinct from genetic interaction- and gene expression-based genetic networks in that they leverage evolutionary, rather than functional, data. Thus, coevolution networks infer functionally conserved relationships among orthologous genes across entire lineages, whereas genetic networks infer functional relationships among genes in a single extant species or strain (irrespective of whether these relationships are conserved in other species or not). Gene coevolution networks are also distinct from networks constructed from correlated presence and absence patterns of orthologs across a lineage (an approach known as phylogenetic profiling (Cokus *et al*. 2007; Pellegrini 2012)) in that coevolutionary networks depict relationships among orthologs conserved in the majority of taxa. Examination of the global coevolution network, communities therein, and signatures of orthologous gene coevolution among bioprocesses, complexes, and pathways reveals that the network reflects the hierarchy of cellular function.

Comparison of the budding yeast coevolution network to the genetic interaction-based network of *S. cerevisiae* revealed numerous notable similarities and differences. For example, both methods found that gene essentiality significantly impacts connectivity wherein essential genes / orthologous genes are more densely connected than nonessential genes / orthologous genes (Fig. 2). This finding suggests that genes with more essential cellular functions are more likely central hubs in the coevolution network (Mnaimneh *et al*. 2004; Costanzo *et al*. 2010, 2016, 2021; Wisecaver *et al*. 2017). Similarities were also observed among genes with broadly conserved functions. For example, the majority of genes / orthologs connected to *CDC6*, a gene required for the fundamental and widely conserved process of DNA replication (Hartwell *et al*. 1970), in the orthologous gene coevolution network and the genetic interaction-based network were the same (Costanzo *et al*. 2010; Usaj *et al*. 2017).

Similarities between genetic interaction and gene coevolution networks were also observed when examining the impact of gene deletion(s) on fitness. For example, in the gene coevolution network, negative fitness outcomes were associated with genetic perturbations of community 1, which is enriched in essential genes, and buffering is largely observed when genes belong to the same community (Figure 4). In the genetic interaction network, deletion of genes from the same essential complex resulted in negative genetic interactions whereas deletion of genes from the same nonessential complex were often associated with positive genetic interactions (Costanzo *et al*. 2016). These striking similarities suggest that, despite using different data types to infer genetic interaction networks and gene coevolutionary networks (i.e., functional and evolutionary data, respectively), functional associations between genes can be encoded in their coevolutionary histories; thus, functional insights can be inferred from gene coevolution networks.

In contrast, differences between the two networks are likely driven by the fact that not all parts of the genetic interaction-based network of any single organism are conserved across an entire lineage (Tong *et al*. 2001; Pan *et al*. 2004; Onge *et al*. 2007; Mani *et al*. 2008; Dixon *et al*. 2008; Lehner 2011; Boucher and Jenna 2013; Lind *et al*. 2015; Sorrells and Johnson 2015; Monaco *et al*. 2015; Yang and Wittkopp 2017). The more distinct the evolutionary histories of genes or pathways of species used to construct an orthologous gene coevolution network, the more divergent the topologies of the genetic interaction-based network of a species in that lineage will be from the coevolution network of the entire lineage. For example, *CKI1*, a choline kinase, gene connectivity substantially differed in the two networks. This may be in part driven by an ancient whole genome duplication event and retention of the duplicate gene copy in some, but not all, budding yeast species (Marcet-Houben and Gabaldón 2015; Wolfe 2015). Taken together, these results indicate that similarities and differences between networks inferred using orthologous gene coevolution from a lineage and networks inferred based on genetic interactions from a single organism are driven by divergence in individual organisms’ genetic networks; thus, these methods offer distinct insights into functional associations among genes.

Another difference between the two networks is that the budding yeast coevolution network offers novel evolutionary insights, which cannot be inferred from genetic interaction networks in a single species. For example, hubs of genes do not only represent functionally related genes but also genes whose function has been maintained across long evolutionary timescales. Furthermore, interpolation of the gene coevolution network and one-dimensional and three-dimensional chromosome structure offers novel insights into the interplay of chromosome structure and coevolution. Despite there being few known examples of inter-chromosomal gene associations (Gibbons *et al*. 2015), we find extensive signatures of inter- and long-range intra-chromosomal coevolution (Fig. 5, S21-S22), which suggests that gene function, not location, drives orthologous gene coevolution over macroevolutionary timescales. These results uncover a previously underappreciated degree of genome-wide coevolution that has been maintained over millions of years of budding yeast evolution, raising the hypothesis that evolution and function of eukaryotic genomes are best viewed as extensively linked ensembles of genes.

In summary, we highlight complementary and novel insights that can be inferred using coevolutionary networks compared to other methods to infer genetic networks. Insights and methods used herein will facilitate the generation, interpretation, and utility of these networks for other lineages in the tree of life.

## Methods

### Inferring gene coevolution

To infer gene coevolution across ∼400 million years of budding yeast evolution, we first obtained 2,408 orthologous sets of genes (hereafter referred to OGs) from 332 species (Shen *et al*. 2018). These 2,408 orthologous genes are from diverse GO bioprocesses but are underrepresented for gene functions known to be present in multiple copies, such as transposons and hexose transporters (Table S5). Thus, we conclude that the 2,408 orthologous sets of genes span a broad range of cellular and molecular functions. Examination of over and underrepresentation of genes from the various chromosomes of *S. cerevisiae* and *C. albicans* revealed no chromosome was over or underrepresented in the 2,408 orthologs (Table S6), suggesting each chromosome is equally represented in our dataset.

Next, we calculated covariation of relative evolutionary rates of all 2,898,028 pairs from the 2,408 orthologous sets of genes. To do so, we developed the CovER (<u>Cov</u>arying <u>E</u>volutionary <u>R</u>ates) pipeline for high-throughput genome-scale analyses of orthologous gene covariation based on the mirror tree principle (Fig. 1). The mirror tree principle is conceptually similar to phylogenetic profiling—wherein correlations in gene presence/absence patterns across a phylogeny are used to identify functionally related genes (Pellegrini *et al*. 1999)—but instead uses correlations in orthologous genes’ relative evolutionary rates (Pazos and Valencia 2001; Clark *et al*. 2012; de Juan *et al*. 2013).

To implement the CovER pipeline, single gene trees constrained to the species topology were first inferred using IQ-TREE, v1.6.11 (Nguyen *et al*. 2015) (Fig. 1). Thereafter, all pairwise combinations of gene trees were examined for significant signatures of coevolution (Fig. 1B). Differences in taxon occupancy between gene trees are accounted for by pruning both phylogenies to the set of maximally shared taxa. To mitigate the influence of factors that can lead to high false positive rates, such as time since speciation and mutation rate, and increase the statistical power of calculating gene coevolution, branch lengths were transformed into relative rates by correcting the gene tree branch length by the corresponding branch length in the species phylogeny (Sato *et al*. 2005; Clark *et al*. 2012; Chikina *et al*. 2016). Single data point outliers (defined as having corrected branch lengths greater than five) are known to cause false positive correlations and were removed (Clark *et al*. 2012). Branch lengths were then Z-transformed and a Pearson correlation coefficient was calculated for each pair of orthologs. The CovER algorithm has been integrated into PhyKIT, a UNIX toolkit for phylogenomic analysis (Steenwyk *et al*. 2021).

### Network construction

Complex interactions between orthologous gene pairs were further examined using a network wherein nodes represent orthologs and edges connect orthologs that are coevolving. Following our previous work (Steenwyk *et al*. 2021), we considered orthologous gene pairs with a covariation coefficient of 0.825 or greater to have a significant signature of coevolution. This threshold resulted in 60,305 / 2,898,026 (2.08%) significant signatures of coevolution (Fig. S1). To explore the impact of our choice of a covariation coefficient threshold, we examined two measures that describe how densely the network is connected: edge density (the proportion of present edges out of all possible edges) and transitivity (ratio of triangles that are connected to triples); as well as two measures that describe how diffuse the network is: mean distance (average path length among pairs of nodes) and diameter (the longest geodesic distance). Across a wide range of thresholds of significant orthologous gene coevolution (Pearson correlation coefficient range of [0.600-0.900] with a step of 0.005), we found that the choice of threshold had little impact on network structure (Fig. S2).

Network substructure is commonly referred to as community structure and describes a set of orthologs that are more densely connected with each other but more sparsely connected with other sets (or communities) of orthologs. To identify the community structure of our global orthologous gene coevolution network, a hierarchical agglomeration algorithm that conducts greedy optimization of modularity was implemented (Newman 2004).

### Enrichment analysis

To determine functional category enrichment among sets of orthologs, gene ontology (GO) enrichment analysis was conducted. To do so, a background set of GO annotations were curated from the 2,408 orthologous genes (Shen *et al*. 2018). Specifically, for an orthologous group of genes, GO associations were mapped from the representative gene from *S. cerevisiae* (Goffeau *et al*. 1996). If an *S. cerevisiae* gene was not present, the annotation from the representative gene from *C. albicans* was chosen (Jones *et al*. 2004). When neither species was represented in an orthologous group, we considered the function of the orthologous group to be uncertain and did not assign a GO term. Significance in functional enrichment was assessed using a Fischer’s exact test with Benjamini Hochberg multi-test correction (α = 0.05) using goatools, v1.0.11 (Klopfenstein *et al*. 2018). GO annotations were obtained from the Gene Ontology Consortium (http://geneontology.org/; release date: 2020-10-09). Higher-order summaries of GO term lists were constructed using GO slim annotations and REVIGO (Supek *et al*. 2011). Over and underrepresentation of essential genes across communities and genes on the various chromosomes were examined using the same approach in R, v4.0.2 (https://cran.r-project.org/).

### Pathway analysis

To examine coevolution between genes in pathways, we first determined the genes belonging to pathways of interest. To do so, we leveraged pathway information in the KEGG database (Kanehisa *et al*. 2016) and the *Saccharomyces* Genome Database (SGD; https://www.yeastgenome.org/). To determine if there are more signatures of coevolution within a pathway than expected by random chance, we conducted permutation tests. The null distribution was generated by randomly shuffling coevolution coefficients across all ∼3 million orthologous gene pairs 10,000 times and then determining the number of coevolving pairs among the pairs of the pathway of interest for each iteration.

### Integrating gene loss information

To estimate the impact of network perturbation, fitness of single-gene deletions and genetic interaction scores inferred from digenic deletions from were combined with information from the orthologous gene coevolution network (Costanzo *et al*. 2010, 2016, 2021; Usaj *et al*. 2017). For example, the relationship between gene-/ortholog-wise community, connectivity, and fitness in diverse environments was evaluated. To determine if genes / orthologs were equally likely to be lost across communities, we examined patterns of gene losses in *Hanseniaspora* spp., which have undergone extensive gene loss compared to other budding yeasts (Steenwyk *et al*. 2019).

### Projecting the network onto genome structure and organization

To gain insight into the relationship between genome structure and the orthologous gene coevolution network, we projected the network onto the complete chromosome genome assemblies of *S. cerevisiae* and *C. albicans* (Goffeau *et al*. 1996; Jones *et al*. 2004; van het Hoog *et al*. 2007; Muzzey *et al*. 2013). Prior to mapping the network onto the genome assemblies, we investigated genome-wide synteny using orthology information from the Candida Gene Order Browser (Fitzpatrick *et al*. 2010). Thereafter, the network was projected onto each genome assembly using Circos, v0.69 (Krzywinski *et al*. 2009). Examination of the distance between coevolving orthologous genes and chromosomal contacts was conducted using a three-dimensional model of the *S. cerevisiae* genome (Duan *et al*. 2010).

## Data Availability

To facilitate other researchers to explore the gene coevolution information, we created a web application, *the budding yeast coevolution network* (https://github.com/JLSteenwyk/budding_yeast_coevolution_network), written in the R programming language (https://cran.r-project.org/). All other supplementary information including single gene phylogenies used to examine coevolution and Pearson covariation coefficients among relative evolutionary rates for all pairwise combinations of orthologous groups of genes will be available on figshare upon publication (doi: 10.6084/m9.figshare.14501964).

## References

Albalat R., and C. Cañestro, 2016 Evolution by gene loss. Nat. Rev. Genet. 17: 379–391. https://doi.org/10.1038/nrg.2016.39

Ayala R., O. Willhoft, R. J. Aramayo, M. Wilkinson, E. A. McCormack, et al., 2018 Structure and regulation of the human INO80–nucleosome complex. Nature 556: 391–395. https://doi.org/10.1038/s41586-018-0021-6

Baryshnikova A., M. Costanzo, C. L. Myers, B. Andrews, and C. Boone, 2013 Genetic Interaction Networks: Toward an Understanding of Heritability. Annu. Rev. Genomics Hum. Genet. 14: 111–133. https://doi.org/10.1146/annurev-genom-082509-141730

Boucher B., and S. Jenna, 2013 Genetic interaction networks: better understand to better predict. Front. Genet. 4. https://doi.org/10.3389/fgene.2013.00290

Brunette G. J., M. A. Jamalruddin, R. A. Baldock, N. L. Clark, and K. A. Bernstein, 2019 Evolution-based screening enables genome-wide prioritization and discovery of DNA repair genes. Proc. Natl. Acad. Sci. 116: 19593–19599. https://doi.org/10.1073/pnas.1906559116

Chibana H., N. Oka, H. Nakayama, T. Aoyama, B. B. Magee, et al., 2005 Sequence Finishing and Gene Mapping for Candida albicans Chromosome 7 and Syntenic Analysis Against the Saccharomyces cerevisiae Genome. Genetics 170: 1525–1537. https://doi.org/10.1534/genetics.104.034652

Chikina M., J. D. Robinson, and N. L. Clark, 2016 Hundreds of Genes Experienced Convergent Shifts in Selective Pressure in Marine Mammals. Mol. Biol. Evol. 33: 2182–2192. https://doi.org/10.1093/molbev/msw112

Ciniawsky S., I. Grimm, D. Saffian, W. Girzalsky, R. Erdmann, et al., 2015 Molecular snapshots of the Pex1/6 AAA+ complex in action. Nat. Commun. 6: 7331. https://doi.org/10.1038/ncomms8331

Clark N. L., E. Alani, and C. F. Aquadro, 2012 Evolutionary rate covariation reveals shared functionality and coexpression of genes. Genome Res. 22: 714–720. https://doi.org/10.1101/gr.132647.111

Cokus S., S. Mizutani, and M. Pellegrini, 2007 An improved method for identifying functionally linked proteins using phylogenetic profiles. BMC Bioinformatics 8: S7. https://doi.org/10.1186/1471-2105-8-S4-S7

Costanzo M., A. Baryshnikova, J. Bellay, Y. Kim, E. D. Spear, et al., 2010 The Genetic Landscape of a Cell. Science (80-.). 327: 425–431. https://doi.org/10.1126/science.1180823

Costanzo M., B. VanderSluis, E. N. Koch, A. Baryshnikova, C. Pons, et al., 2016 A global genetic interaction network maps a wiring diagram of cellular function. Science (80-.). 353: aaf1420–aaf1420. https://doi.org/10.1126/science.aaf1420

Costanzo M., E. Kuzmin, J. van Leeuwen, B. Mair, J. Moffat, et al., 2019 Global Genetic Networks and the Genotype-to-Phenotype Relationship. Cell 177: 85–100. https://doi.org/10.1016/j.cell.2019.01.033

Costanzo M., J. Hou, V. Messier, J. Nelson, M. Rahman, et al., 2021 Environmental robustness of the global yeast genetic interaction network. Science (80-.). 372: eabf8424. https://doi.org/10.1126/science.abf8424

Dixon S. J., Y. Fedyshyn, J. L. Y. Koh, T. S. K. Prasad, C. Chahwan, et al., 2008 Significant conservation of synthetic lethal genetic interaction networks between distantly related eukaryotes. Proc. Natl. Acad. Sci. 105: 16653–16658. https://doi.org/10.1073/pnas.0806261105

Duan Z., M. Andronescu, K. Schutz, S. McIlwain, Y. J. Kim, et al., 2010 A three-dimensional model of the yeast genome. Nature 465: 363–367. https://doi.org/10.1038/nature08973

Dujon B., 2010 Yeast evolutionary genomics. Nat. Rev. Genet. 11: 512–524. https://doi.org/10.1038/nrg2811

Findlay G. D., J. L. Sitnik, W. Wang, C. F. Aquadro, N. L. Clark, et al., 2014 Evolutionary Rate Covariation Identifies New Members of a Protein Network Required for Drosophila melanogaster Female Post-Mating Responses, (J. Zhang, Ed.). PLoS Genet. 10: e1004108. https://doi.org/10.1371/journal.pgen.1004108

Fitzpatrick D. A., P. O’Gaora, K. P. Byrne, and G. Butler, 2010 Analysis of gene evolution and metabolic pathways using the Candida Gene Order Browser. BMC Genomics 11: 290. https://doi.org/10.1186/1471-2164-11-290

GeneOntologyConsortium, 2004 The Gene Ontology (GO) database and informatics resource. Nucleic Acids Res. 32: 258D – 261. https://doi.org/10.1093/nar/gkh036

Gibbons J. G., A. T. Branco, S. Yu, and B. Lemos, 2014 Ribosomal DNA copy number is coupled with gene expression variation and mitochondrial abundance in humans. Nat. Commun. 5: 4850. https://doi.org/10.1038/ncomms5850

Gibbons J. G., A. T. Branco, S. A. Godinho, S. Yu, and B. Lemos, 2015 Concerted copy number variation balances ribosomal DNA dosage in human and mouse genomes. Proc. Natl. Acad. Sci. U. S. A. 112: 2485–2490. https://doi.org/10.1073/pnas.1416878112

Goffeau A., B. G. Barrell, H. Bussey, R. W. Davis, B. Dujon, et al., 1996 Life with 6000 Genes. Science (80-.). 274: 546–567. https://doi.org/10.1126/science.274.5287.546

Goh C.-S., A. A. Bogan, M. Joachimiak, D. Walther, and F. E. Cohen, 2000 Co-evolution of proteins with their interaction partners 11 edited by B. Honig. J. Mol. Biol. 299: 283–293. https://doi.org/10.1006/jmbi.2000.3732

Hartwell L. H., J. Culotti, and B. Reid, 1970 Genetic control of the cell-division cycle in yeast. I. Detection of mutants. Proc. Natl. Acad. Sci. U. S. A. 66: 352–9. https://doi.org/10.1073/pnas.66.2.352

Hoog M. van het, T. J. Rast, M. Martchenko, S. Grindle, D. Dignard, et al., 2007 Assembly of the Candida albicans genome into sixteen supercontigs aligned on the eight chromosomes. Genome Biol. 8: R52. https://doi.org/10.1186/gb-2007-8-4-r52

Hosaka K., T. Kodaki, and S. Yamashita, 1989 Cloning and characterization of the yeast CKI gene encoding choline kinase and its expression in Escherichia coli. J. Biol. Chem. 264: 2053–9.

Huang J.-W., A. Acharya, A. Taglialatela, T. S. Nambiar, R. Cuella-Martin, et al., 2020 MCM8IP activates the MCM8-9 helicase to promote DNA synthesis and homologous recombination upon DNA damage. Nat. Commun. 11: 2948. https://doi.org/10.1038/s41467-020-16718-3

Hurst L. D., C. Pál, and M. J. Lercher, 2004 The evolutionary dynamics of eukaryotic gene order. Nat. Rev. Genet. 5: 299–310. https://doi.org/10.1038/nrg1319

Jones T., N. A. Federspiel, H. Chibana, J. Dungan, S. Kalman, et al., 2004 The diploid genome sequence of Candida albicans. Proc. Natl. Acad. Sci. 101: 7329–7334. https://doi.org/10.1073/pnas.0401648101

Juan D. de, F. Pazos, and A. Valencia, 2013 Emerging methods in protein co-evolution. Nat. Rev. Genet. 14: 249–261. https://doi.org/10.1038/nrg3414

Kanehisa M., Y. Sato, M. Kawashima, M. Furumichi, and M. Tanabe, 2016 KEGG as a reference resource for gene and protein annotation. Nucleic Acids Res. 44: D457–D462. https://doi.org/10.1093/nar/gkv1070

Kim K., K. H. Kim, M. K. Storey, D. R. Voelker, and G. M. Carman, 1999 Isolation and characterization of the Saccharomyces cerevisiae EKI1 gene encoding ethanolamine kinase. J. Biol. Chem. 274: 14857–66. https://doi.org/10.1074/jbc.274.21.14857

Kim W. K., D. M. Bolser, and J. H. Park, 2004 Large-scale co-evolution analysis of protein structural interlogues using the global protein structural interactome map (PSIMAP). Bioinformatics 20: 1138–1150. https://doi.org/10.1093/bioinformatics/bth053

Klopfenstein D. V., L. Zhang, B. S. Pedersen, F. Ramírez, A. Warwick Vesztrocy, et al., 2018 GOATOOLS: A Python library for Gene Ontology analyses. Sci. Rep. 8: 10872. https://doi.org/10.1038/s41598-018-28948-z

Krzywinski M., J. Schein, I. Birol, J. Connors, R. Gascoyne, et al., 2009 Circos: An information aesthetic for comparative genomics. Genome Res. 19: 1639–1645. https://doi.org/10.1101/gr.092759.109

Kuzmin E., B. VanderSluis, W. Wang, G. Tan, R. Deshpande, et al., 2018 Systematic analysis of complex genetic interactions. Science (80-.). 360: eaao1729. https://doi.org/10.1126/science.aao1729

Lehner B., 2011 Molecular mechanisms of epistasis within and between genes. Trends Genet. 27: 323–331. https://doi.org/10.1016/j.tig.2011.05.007

Lezon T. R., J. R. Banavar, M. Cieplak, A. Maritan, and N. V. Fedoroff, 2006 Using the principle of entropy maximization to infer genetic interaction networks from gene expression patterns. Proc. Natl. Acad. Sci. 103: 19033–19038. https://doi.org/10.1073/pnas.0609152103

Lind A. L., J. H. Wisecaver, T. D. Smith, X. Feng, A. M. Calvo, et al., 2015 Examining the evolution of the regulatory circuit controlling secondary metabolism and development in the fungal genus Aspergillus. PLoS Genet. 11: e1005096. https://doi.org/10.1371/journal.pgen.1005096

Mani R., R. P. St.Onge, J. L. Hartman, G. Giaever, and F. P. Roth, 2008 Defining genetic interaction. Proc. Natl. Acad. Sci. 105: 3461–3466. https://doi.org/10.1073/pnas.0712255105

Marcet-Houben M., and T. Gabaldón, 2015 Beyond the Whole-Genome Duplication: Phylogenetic Evidence for an Ancient Interspecies Hybridization in the Baker’s Yeast Lineage, (L. D. Hurst, Ed.). PLOS Biol. 13: e1002220. https://doi.org/10.1371/journal.pbio.1002220

Mixão V., and T. Gabaldón, 2020 Genomic evidence for a hybrid origin of the yeast opportunistic pathogen Candida albicans. BMC Biol. 18: 48. https://doi.org/10.1186/s12915-020-00776-6

Mnaimneh S., A. P. Davierwala, J. Haynes, J. Moffat, W.-T. Peng, et al., 2004 Exploration of Essential Gene Functions via Titratable Promoter Alleles. Cell 118: 31–44. https://doi.org/10.1016/j.cell.2004.06.013

Monaco G., S. van Dam, J. L. Casal Novo Ribeiro, A. Larbi, and J. P. de Magalhães, 2015 A comparison of human and mouse gene co-expression networks reveals conservation and divergence at the tissue, pathway and disease levels. BMC Evol. Biol. 15: 259. https://doi.org/10.1186/s12862-015-0534-7

Muzzey D., K. Schwartz, J. S. Weissman, and G. Sherlock, 2013 Assembly of a phased diploid Candida albicans genome facilitates allele-specific measurements and provides a simple model for repeat and indel structure. Genome Biol. 14: R97. https://doi.org/10.1186/gb-2013-14-9-r97

Newman M. E. J., 2004 Fast algorithm for detecting community structure in networks. Phys. Rev. E 69: 066133. https://doi.org/10.1103/PhysRevE.69.066133

Nguyen L.-T., H. A. Schmidt, A. von Haeseler, and B. Q. Minh, 2015 IQ-TREE: A Fast and Effective Stochastic Algorithm for Estimating Maximum-Likelihood Phylogenies. Mol. Biol. Evol. 32: 268–274. https://doi.org/10.1093/molbev/msu300

Onge R. P. S., R. Mani, J. Oh, M. Proctor, E. Fung, et al., 2007 Systematic pathway analysis using high-resolution fitness profiling of combinatorial gene deletions. Nat. Genet. 39: 199– 206. https://doi.org/10.1038/ng1948

Pan X., D. S. Yuan, D. Xiang, X. Wang, S. Sookhai-Mahadeo, et al., 2004 A Robust Toolkit for Functional Profiling of the Yeast Genome. Mol. Cell 16: 487–496. https://doi.org/10.1016/j.molcel.2004.09.035

Pazos F., and A. Valencia, 2001 Similarity of phylogenetic trees as indicator of protein–protein interaction. Protein Eng. Des. Sel. 14: 609–614. https://doi.org/10.1093/protein/14.9.609

Pellegrini M., E. M. Marcotte, M. J. Thompson, D. Eisenberg, and T. O. Yeates, 1999 Assigning protein functions by comparative genome analysis: Protein phylogenetic profiles. Proc. Natl. Acad. Sci. 96: 4285–4288. https://doi.org/10.1073/pnas.96.8.4285

Pellegrini M., 2012 Using Phylogenetic Profiles to Predict Functional Relationships, pp. 167– 177 in.

Raza Q., J. Y. Choi, Y. Li, R. M. O’Dowd, S. C. Watkins, et al., 2019 Evolutionary rate covariation analysis of E-cadherin identifies Raskol as a regulator of cell adhesion and actin dynamics in Drosophila, (C. Desplan, Ed.). PLOS Genet. 15: e1007720. https://doi.org/10.1371/journal.pgen.1007720

Reuber B. E., E. Germain-Lee, C. S. Collins, J. C. Morrell, R. Ameritunga, et al., 1997 Mutations in PEX1 are the most common cause of peroxisome biogenesis disorders. Nat. Genet. 17: 445–448. https://doi.org/10.1038/ng1297-445

Rokas A., J. H. Wisecaver, and A. L. Lind, 2018 The birth, evolution and death of metabolic gene clusters in fungi. Nat. Rev. Microbiol. https://doi.org/10.1038/s41579-018-0075-3

Sato T., Y. Yamanishi, M. Kanehisa, and H. Toh, 2005 The inference of protein-protein interactions by co-evolutionary analysis is improved by excluding the information about the phylogenetic relationships. Bioinformatics 21: 3482–3489. https://doi.org/10.1093/bioinformatics/bti564

Segal E. S., V. Gritsenko, A. Levitan, B. Yadav, N. Dror, et al., 2018 Gene Essentiality Analyzed by In Vivo Transposon Mutagenesis and Machine Learning in a Stable Haploid Isolate of Candida albicans, (A. Di Pietro, Ed.). MBio 9. https://doi.org/10.1128/mBio.02048-18

Seoighe C., N. Federspiel, T. Jones, N. Hansen, V. Bivolarovic, et al., 2000 Prevalence of small inversions in yeast gene order evolution. Proc. Natl. Acad. Sci. 97: 14433–14437. https://doi.org/10.1073/pnas.240462997

Shen X.-X., D. A. Opulente, J. Kominek, X. Zhou, J. L. Steenwyk, et al., 2018 Tempo and Mode of Genome Evolution in the Budding Yeast Subphylum. Cell 175: 1533-1545.e20. https://doi.org/10.1016/j.cell.2018.10.023

Sorrells T. R., and A. D. Johnson, 2015 Making Sense of Transcription Networks. Cell 161: 714–723. https://doi.org/10.1016/j.cell.2015.04.014

Steenwyk J. L., D. A. Opulente, J. Kominek, X.-X. Shen, X. Zhou, et al., 2019 Extensive loss of cell-cycle and DNA repair genes in an ancient lineage of bipolar budding yeasts, (S. Kamoun, Ed.). PLOS Biol. 17: e3000255. https://doi.org/10.1371/journal.pbio.3000255

Steenwyk J. L., T. J. Buida, A. L. Labella, Y. Li, X.-X. Shen, et al., 2021 PhyKIT: a broadly applicable UNIX shell toolkit for processing and analyzing phylogenomic data., (R. Schwartz, Ed.). Bioinformatics. https://doi.org/10.1093/bioinformatics/btab096

Supek F., M. Bosnjak, N. Skunca, and T. Smuc, 2011 REVIGO summarizes and visualizes long lists of gene ontology terms. PLoS One 6: e21800. https://doi.org/10.1371/journal.pone.0021800

Talsness D. M., K. G. Owings, E. Coelho, G. Mercenne, J. M. Pleinis, et al., 2020 A Drosophila screen identifies NKCC1 as a modifier of NGLY1 deficiency. Elife 9. https://doi.org/10.7554/eLife.57831

Tong A. H. Y., M. Evangelista, A. B. Parsons, H. Xu, G. D. Bader, et al., 2001 Systematic Genetic Analysis with Ordered Arrays of Yeast Deletion Mutants. Science (80-.). 294: 2364–2368. https://doi.org/10.1126/science.1065810

Usaj M., Y. Tan, W. Wang, B. VanderSluis, A. Zou, et al., 2017 TheCellMap.org: A Web-Accessible Database for Visualizing and Mining the Global Yeast Genetic Interaction Network. G3&#58; Genes|Genomes|Genetics 7: 1539–1549. https://doi.org/10.1534/g3.117.040220

Winzeler E. A., D. D. Shoemaker, A. Astromoff, H. Liang, K. Anderson, et al., 1999 Functional characterization of the S-cerevisiae genome by gene deletion and parallel analysis. Science (80-.). 285: 901–906. https://doi.org/10.1126/science.285.5429.901

Wisecaver J. H., A. T. Borowsky, V. Tzin, G. Jander, D. J. Kliebenstein, et al., 2017 A Global Coexpression Network Approach for Connecting Genes to Specialized Metabolic Pathways in Plants. Plant Cell 29: 944–959. https://doi.org/10.1105/tpc.17.00009

Wolfe K. H., 2006 Comparative genomics and genome evolution in yeasts. Philos. Trans. R. Soc. B Biol. Sci. 361: 403–412. https://doi.org/10.1098/rstb.2005.1799

Wolfe K. H., 2015 Origin of the Yeast Whole-Genome Duplication. PLOS Biol. 13: e1002221. https://doi.org/10.1371/journal.pbio.1002221

Yang B., and P. J. Wittkopp, 2017 Structure of the Transcriptional Regulatory Network Correlates with Regulatory Divergence in Drosophila. Mol. Biol. Evol. 34: 1352–1362. https://doi.org/10.1093/molbev/msx068

