## Supplementary Figures for "An orthologous gene coevolution network provides insight into eukaryotic cellular and genomic structure and function"

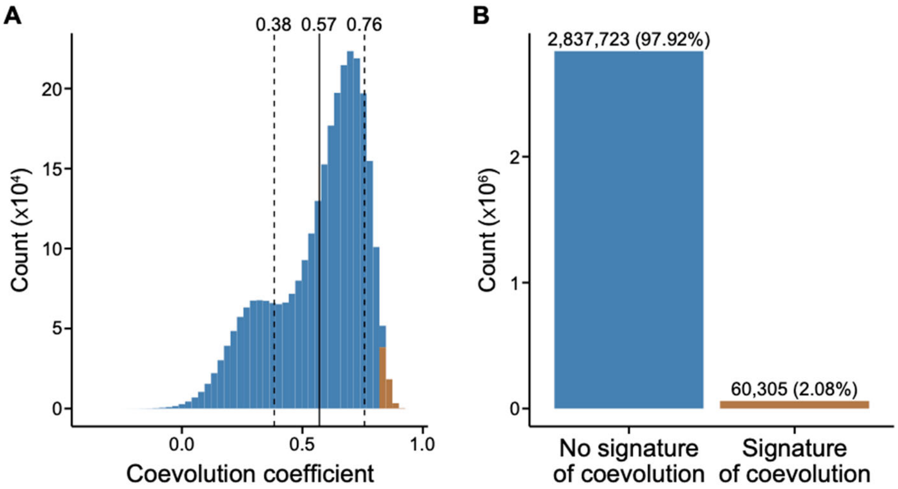


**Figure S1. A conservative threshold for signatures of gene coevolution was used to construct the gene coevolution network.** (A) Distribution of coevolution coefficients. The average coevolution coefficient is 0.57 and is represented by a solid line. One standard deviation away from the average is depicted using a dashed line. Insignificant coevolution coefficients are shown in blue whereas significant coevolution coefficients, which are defined as having values greater than or equal to 0.825, are shown in gold. (B) There are 60,305 significant signatures of gene coevolution whereas there are 2,837,723 insignificant signatures of gene coevolution.

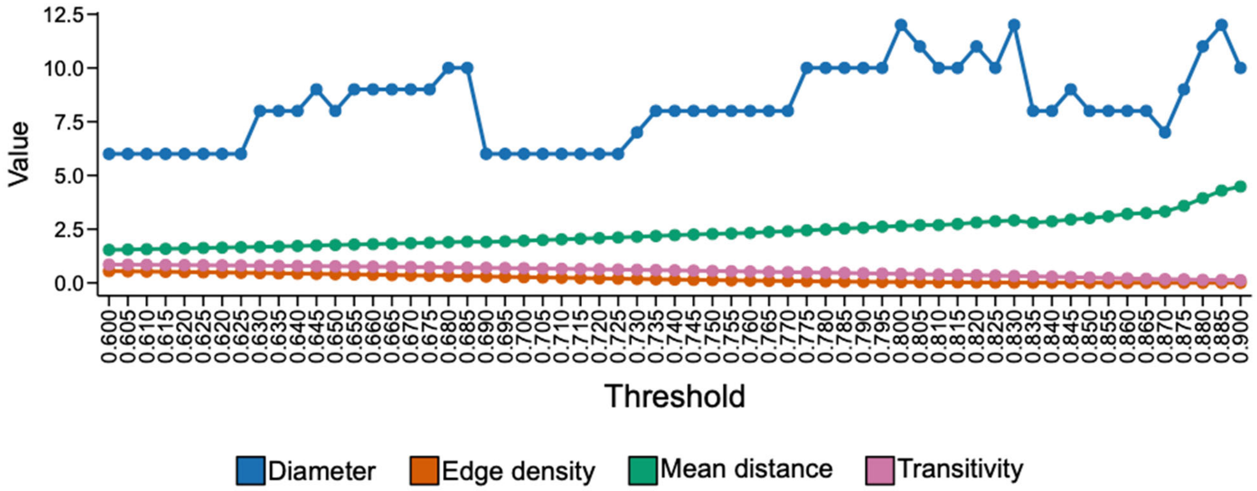


**Figure S2. Various thresholds of significant gene coevolution had little impact on overall network features.** To determine the impact of various thresholds of coevolution coefficients on the resulting network, we examined network diameter, edge density, mean distance, and transitivity. Network properties were similar regardless of the threshold used.

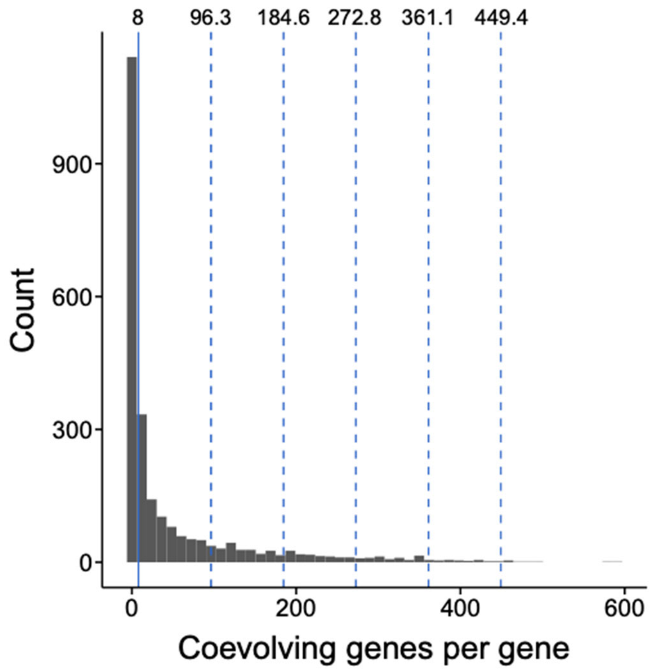


**Figure S3. Distribution of node degrees.** The median number of node degrees is eight (solid line). The dashed lines represent the median plus one, two, three, four, or five standard deviations.

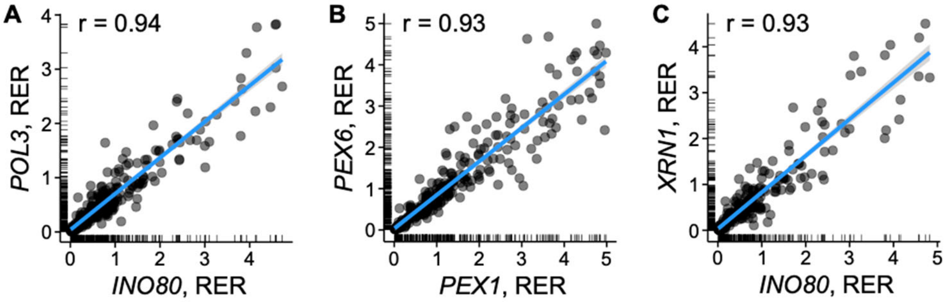


**Figure S4. Three gene pairs with the strongest signatures of gene coevolution.** (A) *INO80*, a gene that encodes a nucleosome spacing factor, and *POL3*, a gene that encodes the catalytic subunit of DNA polymerase delta, are significantly coevolving. *PEX1* and *PEX6*, which form a heterodimer, are also coevolving. Similarly, *INO80* and *XRN1*, which encodes an exoribonuclease, have significant signatures of evolution. Each dot represents a branch in the gene tree phylogeny.

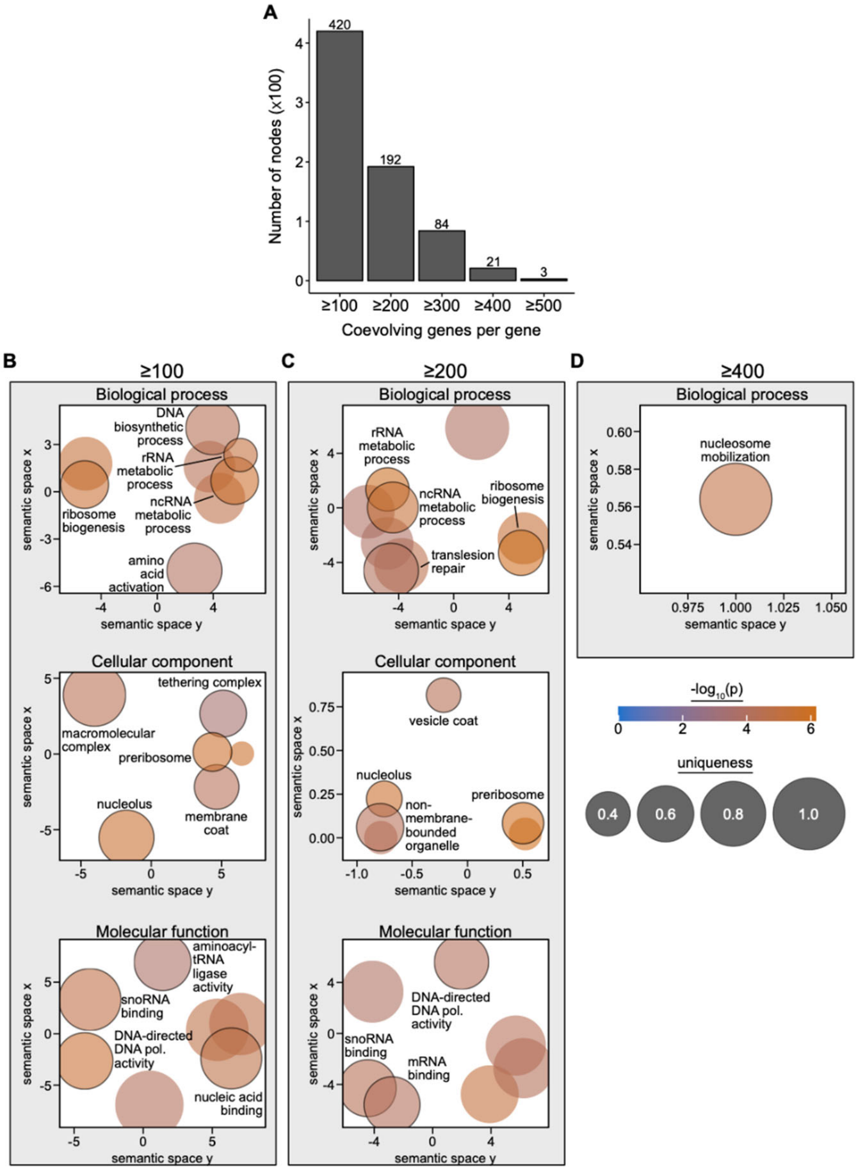


**Figure S5. Gene ontology enrichment of genes with high degrees reveals functional categories associated with highlight coordinated processes.** To determine what functional categories of genes are densely connected to other genes, we conducted GO enrichment analysis. (A) To do so, we first binned genes into groups having ≥100, ≥200, ≥300, ≥400, and ≥500 coevolving genes per gene. (B, C, and D) Enriched terms were observed among genes coevolving with ≥100, ≥200, and ≥400 genes. Enriched terms among genes coevolving with ≥100 and ≥200 genes included those involved in ribosome biogenesis and processes involving nucleic acids such as their binding and synthesis. (D) Among genes coevolving with ≥400 genes, there was one enriched term, nucleosome mobilization, which is associated with chromatin remodeling. In B-D, significantly enriched terms ($\alpha = 0.05$) are represented as circles in semantic space. Uniqueness, a measure of GO term dissimilarity to all other enriched terms, is represented by circle size. Circle color is representative of $-\log_{10}$ transformed p-value. Highlighted enriched terms are written within the figure and their corresponding circle has a black outline. Complete GO enrichment analyses are reported in Table S1.

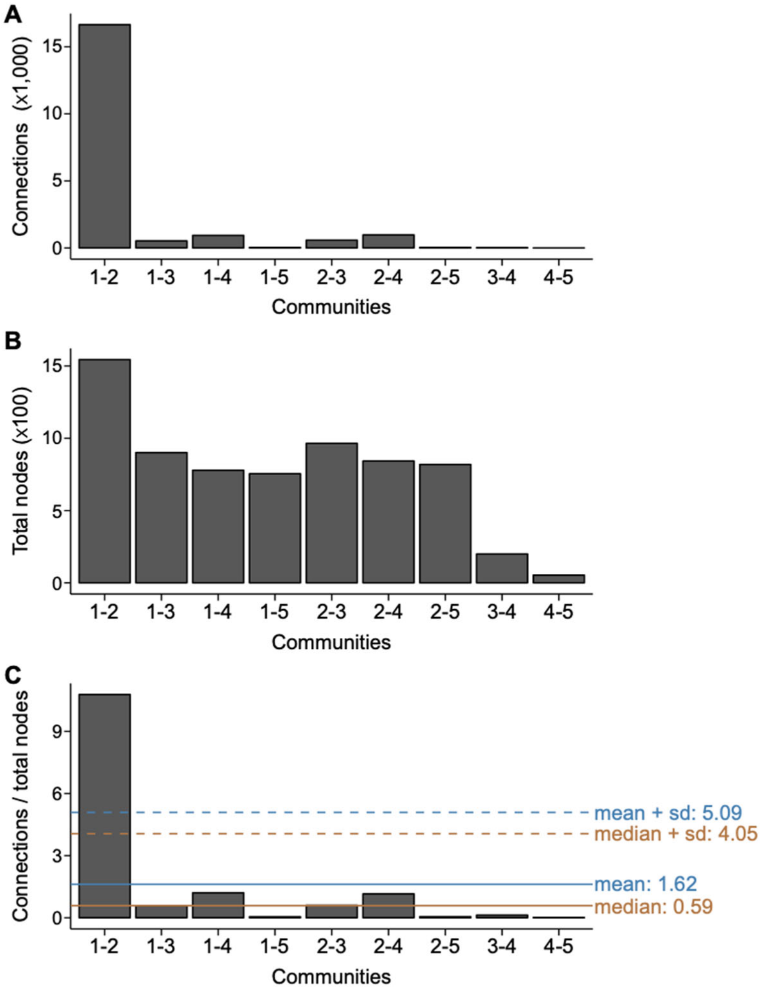


**Figure S6. Communities one and two are highly connected.** (A) Examination of the total number of connections between communities reveals community one and two are highly connected. Correcting the number of connections between communities by (B) the total number of genes in each community reveals that (C) communities one and two are exceptionally interconnected. Mean and median connections between communities corrected by the total number of genes in each community is represented in a blue and orange solid line, respectively. The mean or median plus one standard deviation is shown in a dashed line.

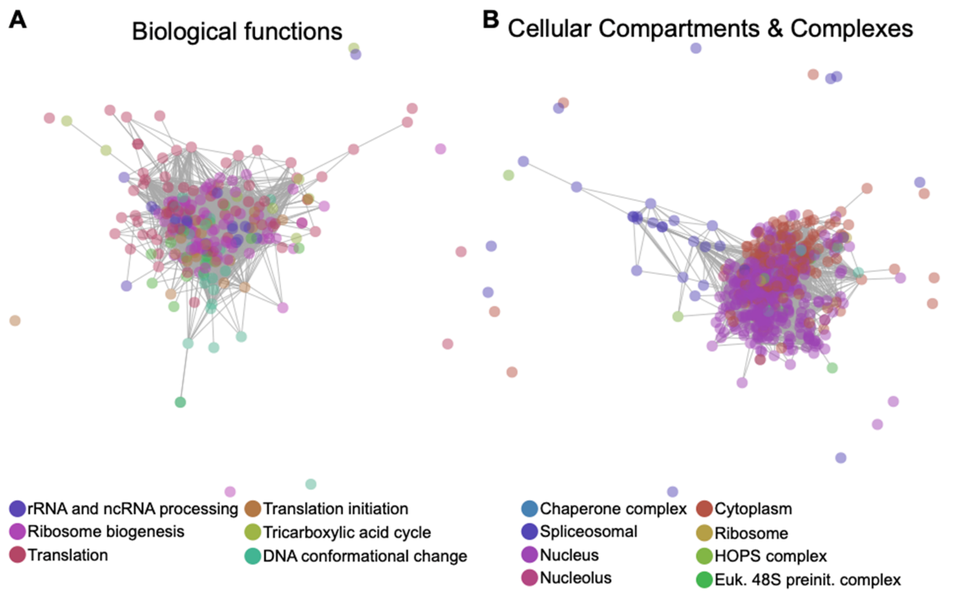


**Figure S7. Subnetworks of broad categories reveal connections and bridges between biological functions and cellular compartments and complexes.** (A) Diverse biological functions derived from enriched terms across communities reveal a high degree of interconnectedness. (B) Examination of cellular compartments and complexes uncovers cellular structure. For example, the nucleus (purple) and cytoplasm (red) are adjacent and intertwined in the network and are bridged by genes that perform spliceosome-related functions (dark blue). Similarly, transcripts are created and processed by the spliceosome in the nucleus and before being transported to the cytoplasm for translation.

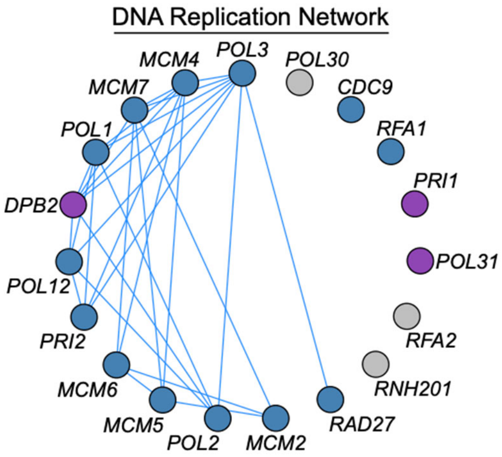


**Figure S8. DNA replication as an exemplary pathway with signatures of gene-gene coevolution.** Gene-gene coevolution network among genes involved in DNA replication. Nodes represent genes and edges connect coevolving genes. Genes are arranged counter-clockwise according to decreasing numbers of degrees, or coevolving genes per gene, in the subnetwork.

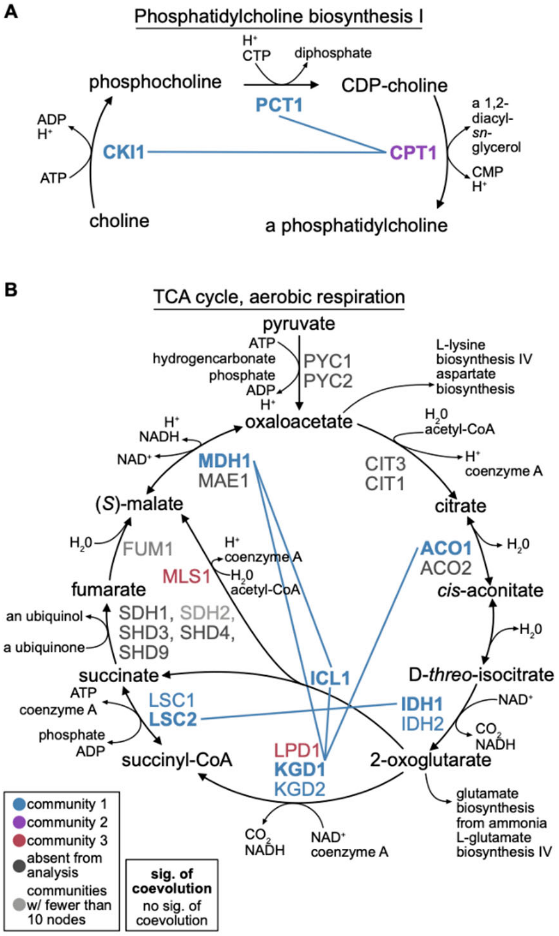


**Figure S9. Exemplary pathways that are coevolving.** (A) Phosphatidylcholine, the major phospholipid in organelle membranes, is synthesized by the genes *CKI1*, *PCT1*, and *CPT1*. *CPT1* is coevolving with *CKI1* and *PCT1*. (B) The tricarboxylic acid cycle (TCA cycle; also referred to as the Krebs cycle or citric acid cycle) is a key component of aerobic respiration in cells. Many genes in the TCA cycle are coevolving with one another.

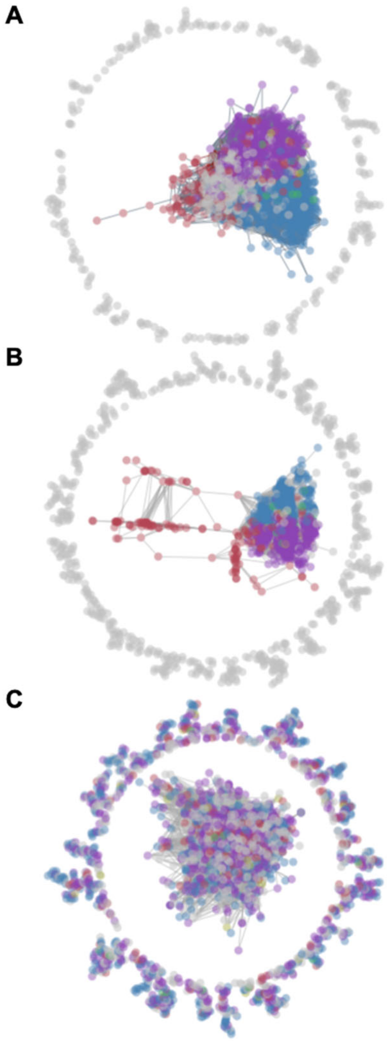


**Figure S10. The coevolution genetic network and genetic interaction network differ.** (A) Edges identified in the coevolution genetic network and genetic interaction network are combined into one super network. (B) The global network inferred using gene-gene coevolution. (C) The genetic interaction network among only significant genetic interaction scores. Nodes are genes and edges connect genes with significant signatures of coevolution or genetic interactions. Genes in community one, two, three, four, five, and all other small communities are depicted in blue, purple, red, yellow, green, and grey respectively.

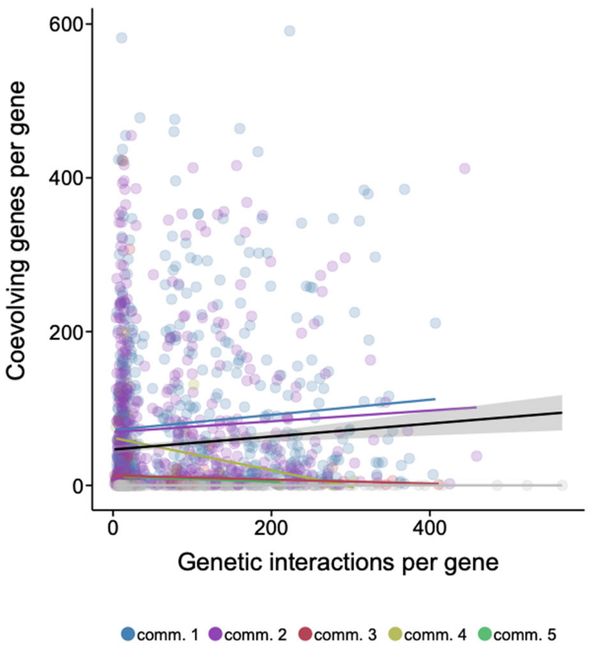


**Figure S11. The coevolution genetic network and genetic interaction network differ in gene connectivity.** Degrees in the coevolutionary network and the genetic interaction network are not related to one another. Each circle represents a gene and their color reflects the community they belong to. Lines represent linear model regressions between the degrees of each network and their color reflects the community they represent; a black line with a 95% confidence in grey represents the linear model regression across all communities.

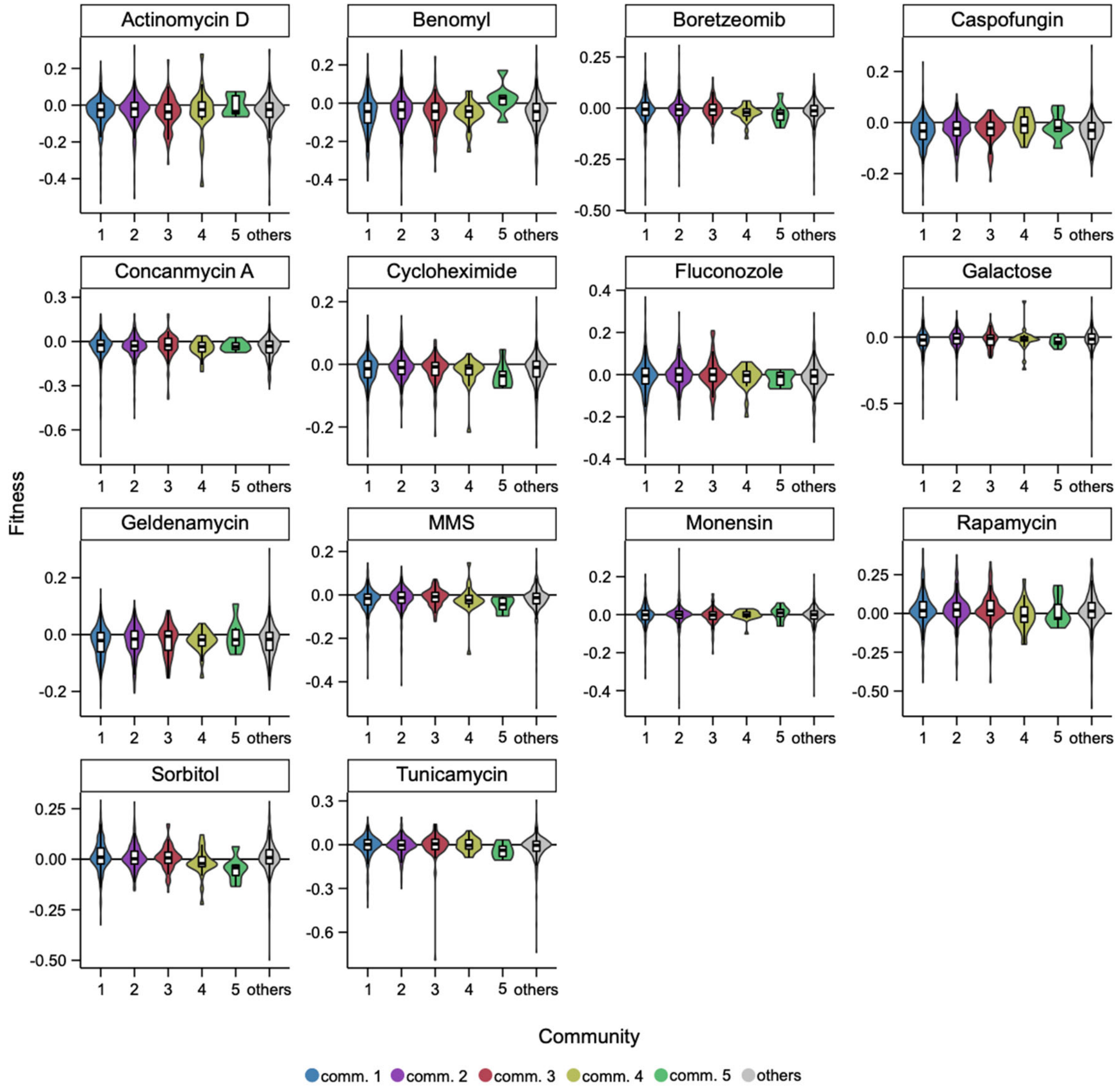


**Figure S12. Community-dependent variation in fitness of single-gene knockouts across environments.** Fitness (y-axis) among single-gene knockouts is in part dependent on the community (x-axis) to which the gene belongs to across 14 environments.

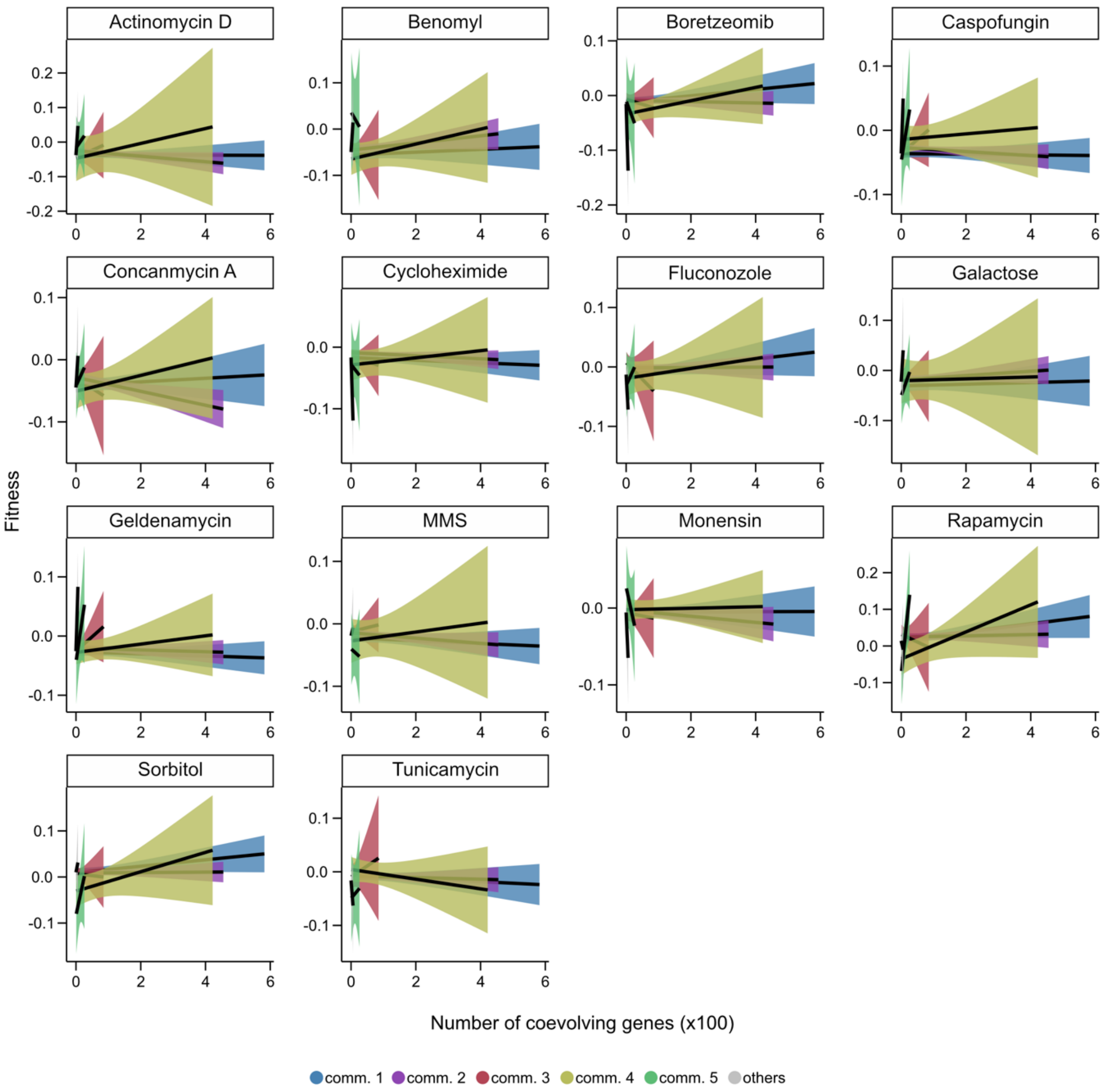


**Figure S13. Community-dependent variation in fitness of single-gene knockouts is in part dependent on gene connectivity across environments.** Fitness (y-axis) among single-gene knockouts is in part dependent on the community to which the gene belongs to across 14 environments as well as gene connectivity as measured by the number of coevolving genes per gene (x-axis).

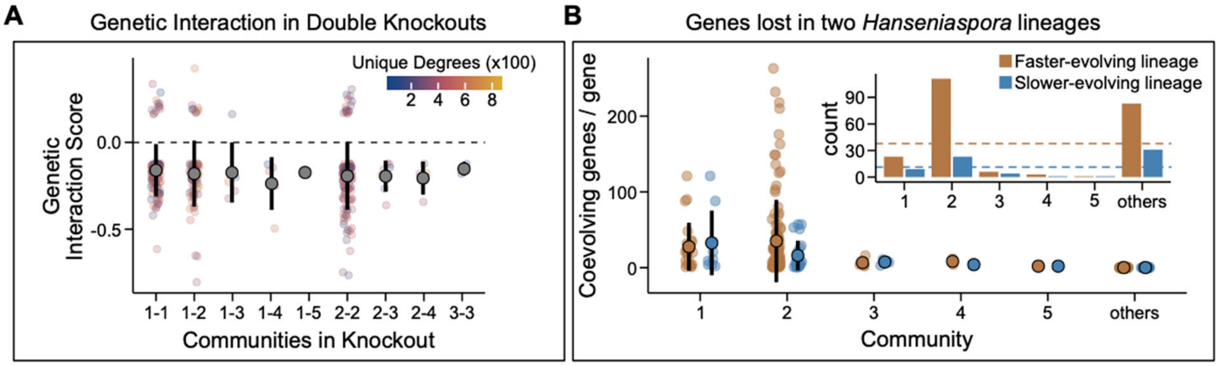


**Figure S14. Digenic gene losses greatly impact cellular fitness and genes lost in the yeast genus *Hanseniaspora* are lost asymmetrically across communities.** (A) Negative genetic interaction scores reflecting a worse phenotypic impact when two genes are deleted are more frequently observed across double knockouts. Genetic interactions scores between different community combinations represented in the double knockout were not significantly different (p-value > 0.05; Kruskal-Wallis rank sum test). (B) Gene loss occurs asymmetrically across communities in two *Hanseniaspora* lineages. The inset depicts total counts of losses per community and dashed lines reflect average losses for each lineage.

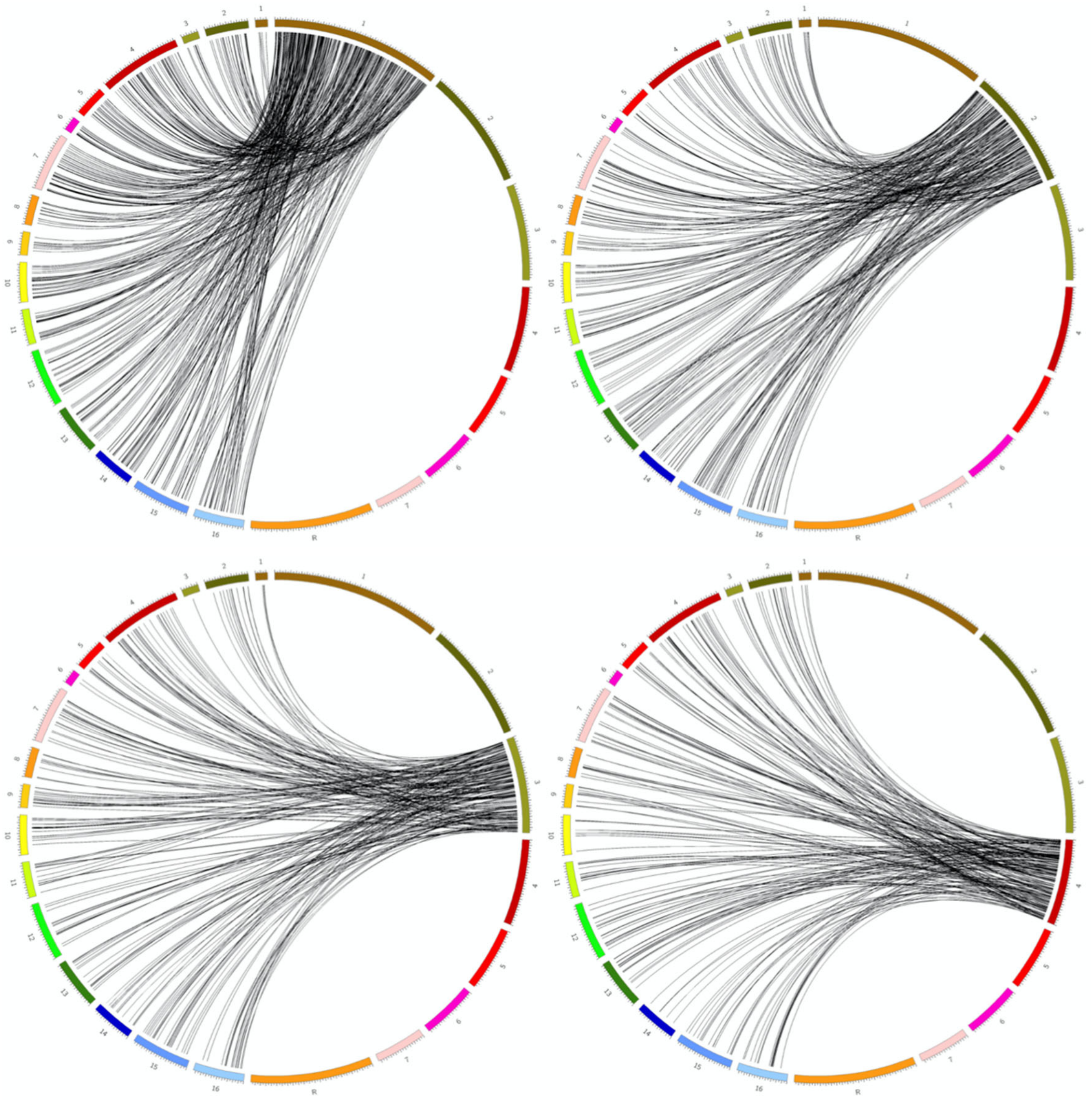

**Figure S15. Lack of synteny between the *Candida albicans* chromosomes 1, 2, 3, and 4 and *Saccharomyces cerevisiae* chromosomes.** The left 16 chromosomes are the *S. cerevisiae* chromosomes. The right eight chromosomes are the *C. albicans* chromosomes. Links start from a specific *C. albicans* chromosome and are connected to their orthologous gene in the *S. cerevisiae* genome. Gene orthology reveals a lack of synteny between the two genomes. Gene orthology information was obtained from the *Candida* genome browser http://www.candidagenome.org/download/homology/orthologs/C_albicans_SC5314_S_cerevisiae_by_CGOB/.

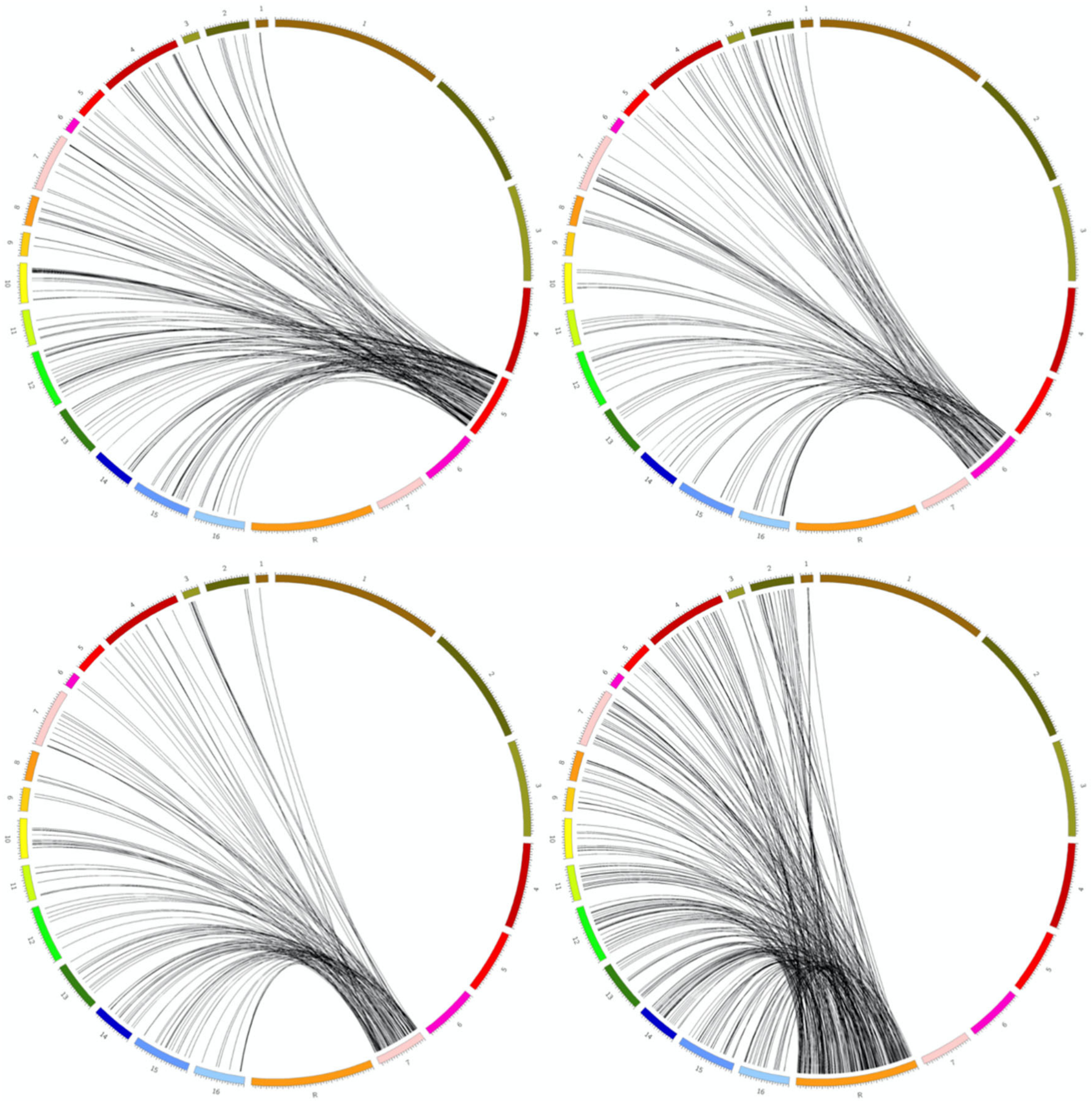

**Figure S16. Lack of synteny between the *Candida albicans* chromosomes 5, 6, 7, and R and *Saccharomyces cerevisiae* chromosomes.** The left 16 chromosomes are the *S. cerevisiae* chromosomes. The right eight chromosomes are the *C. albicans* chromosomes. Links start from a specific *C. albicans* chromosome and are connected to their orthologous gene in the *S. cerevisiae* genome. Gene orthology reveals a lack of synteny between the two genomes Gene orthology information was obtained from the *Candida* genome browser http://www.candidagenome.org/download/homology/orthologs/C_albicans_SC5314_S_cerevisiae_by_CGOB/.

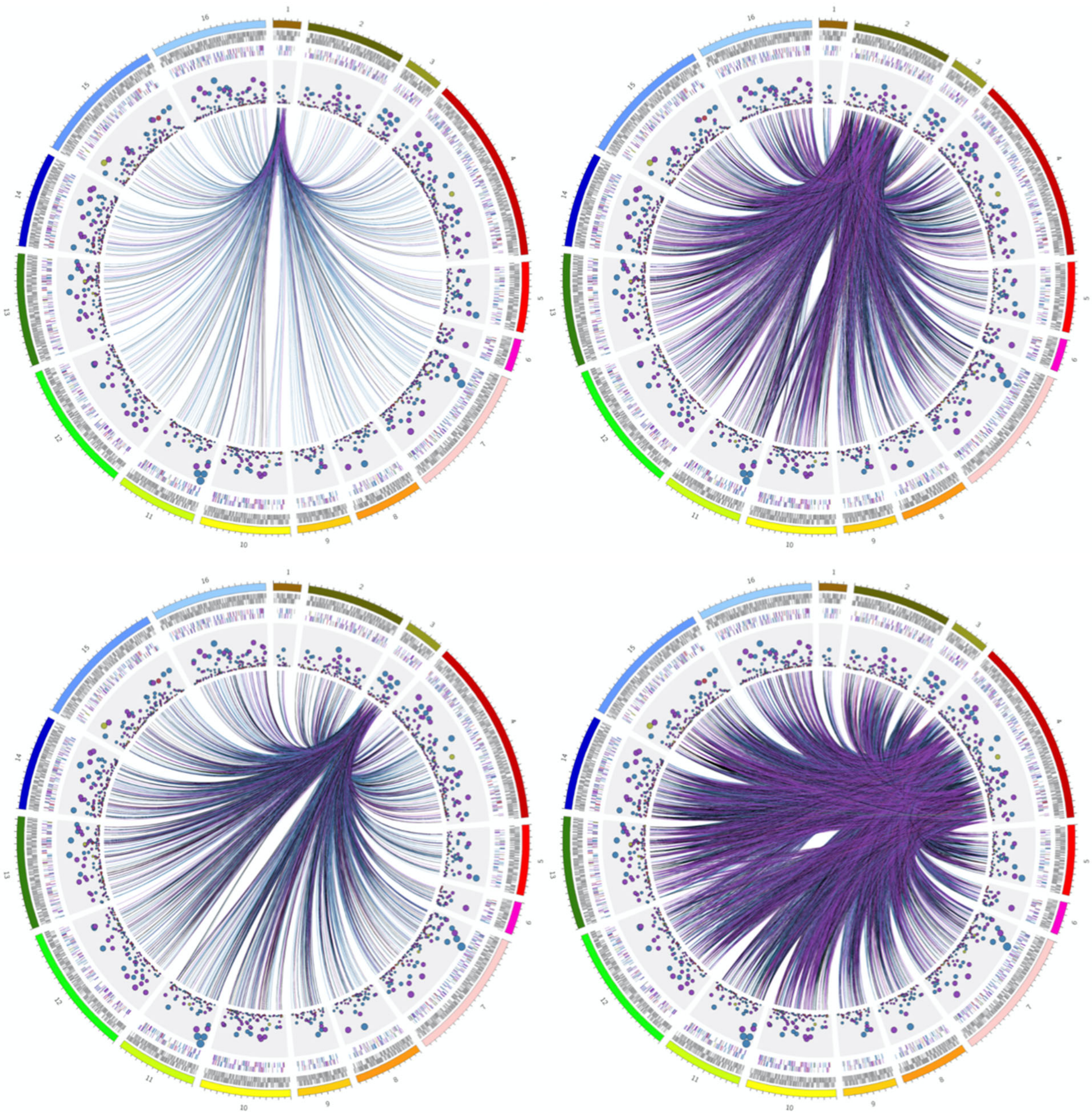

**Figure S17. Genes on chromosomes one through four are coevolving with genes on all other chromosomes in *Saccharomyces cerevisiae*.** The first track depicts the 16 chromosomes of *S. cerevisiae*. The next track depicts the location of all genes on the plus strand followed by the minus strand. The next tracks depict genes present in the dataset and are colored according to the community they belong. The scatter plot depicts the number of coevolving genes a gene has and the larger the circle represents more coevolving genes. The last track depicts links between genes on a highlighted chromosome that are coevolving with other genes across all chromosomes. Plots are arranged in ascending chromosome order from top left to bottom right.

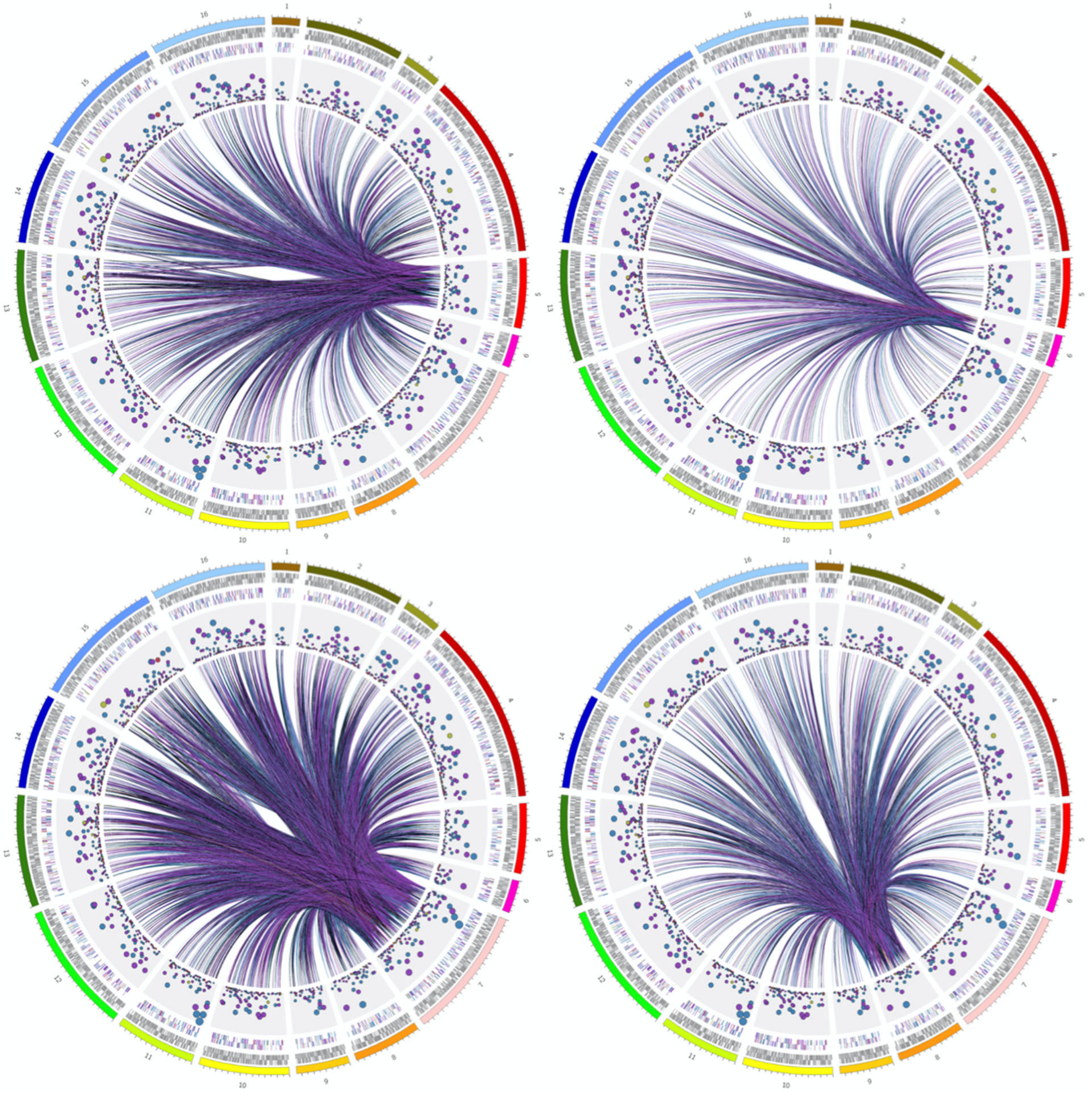

**Figure S18. Genes on chromosomes five through eight are coevolving with genes on all other chromosomes in *Saccharomyces cerevisiae*.** The first six tracks depict the same data as in Figure *N* (Genes on chromosomes one through four are coevolving with genes on all other chromosomes). Links depict genes that are coevolving with genes on a highlighted chromosome. Plots are arranged in ascending chromosome order from top left to bottom right.

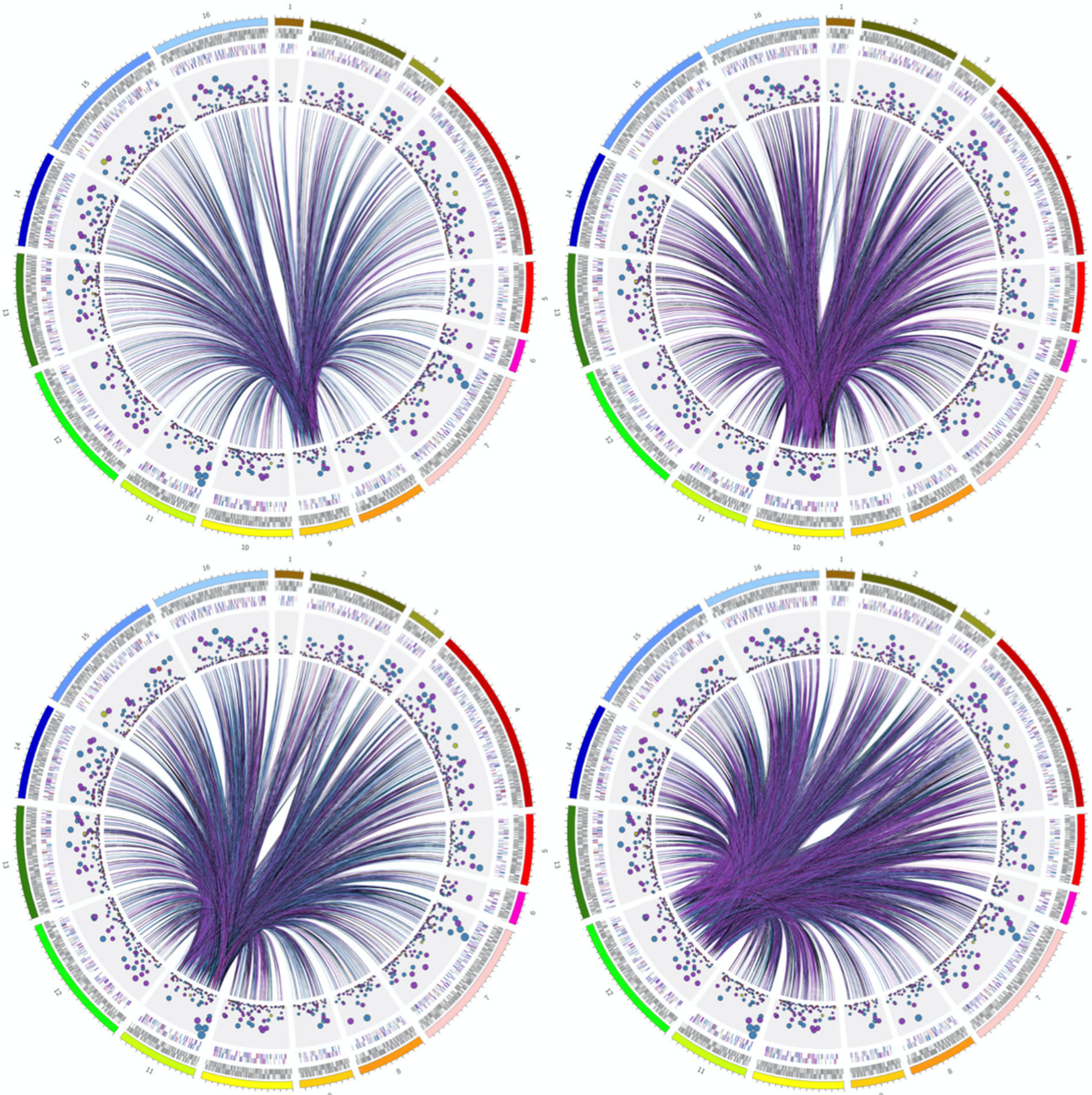

**Figure S19. Genes on chromosomes nine through twelve are coevolving with genes on all other chromosomes in *Saccharomyces cerevisiae*.** The first six tracks depict the same data as in Figure *N* (Genes on chromosomes one through four are coevolving with genes on all other chromosomes). Links depict genes that are coevolving with genes on a highlighted chromosome. Plots are arranged in ascending chromosome order from top left to bottom right.

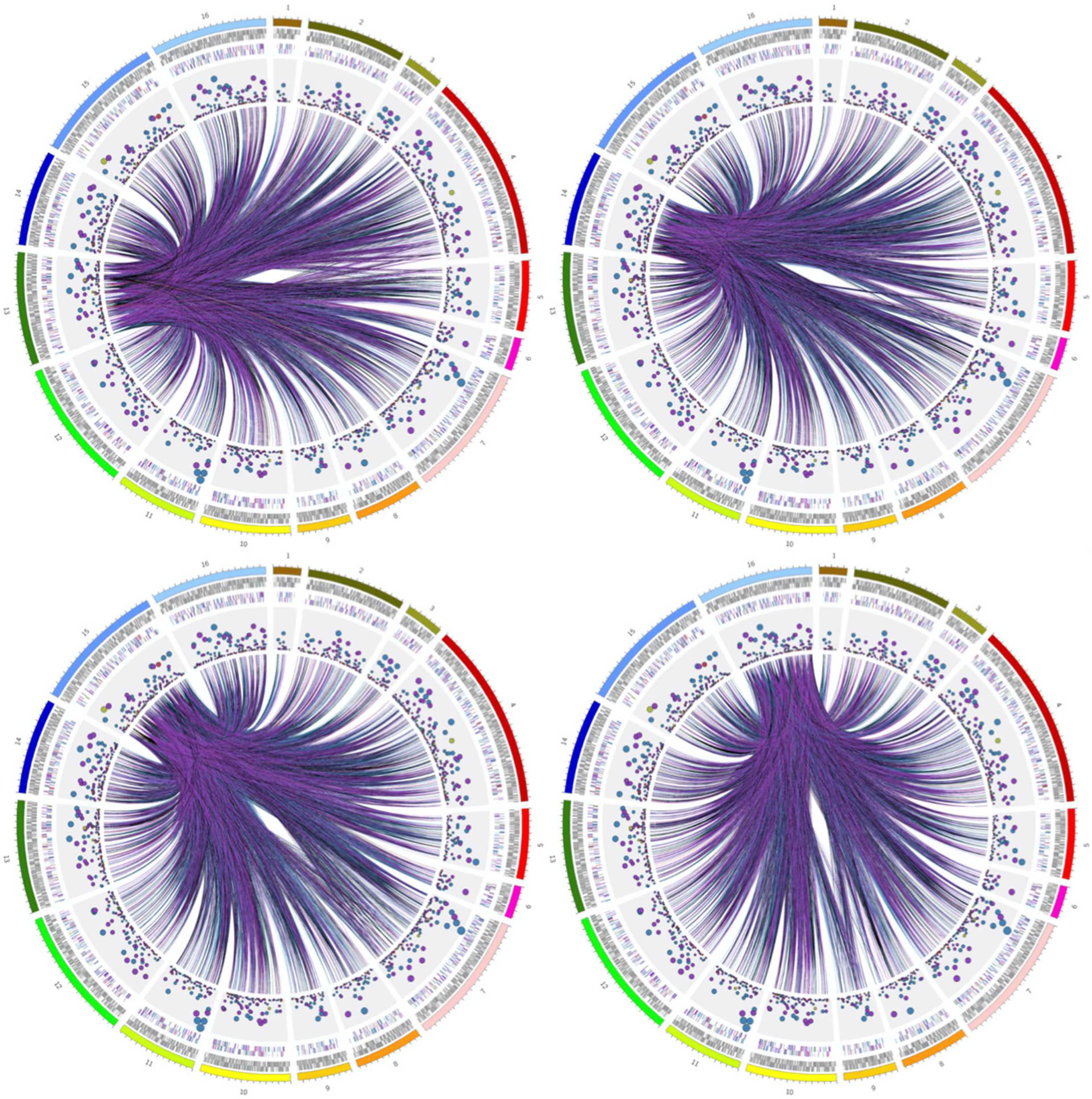

**Figure S20. Genes on chromosomes thirteen through sixteen are coevolving with genes on all other chromosomes in *Saccharomyces cerevisiae*.** The first six tracks depict the same data as in Figure *N* (Genes on chromosomes one through four are coevolving with genes on all other chromosomes). Links depict genes that are coevolving with genes on a highlighted chromosome. Plots are arranged in ascending chromosome order from top left to bottom right.

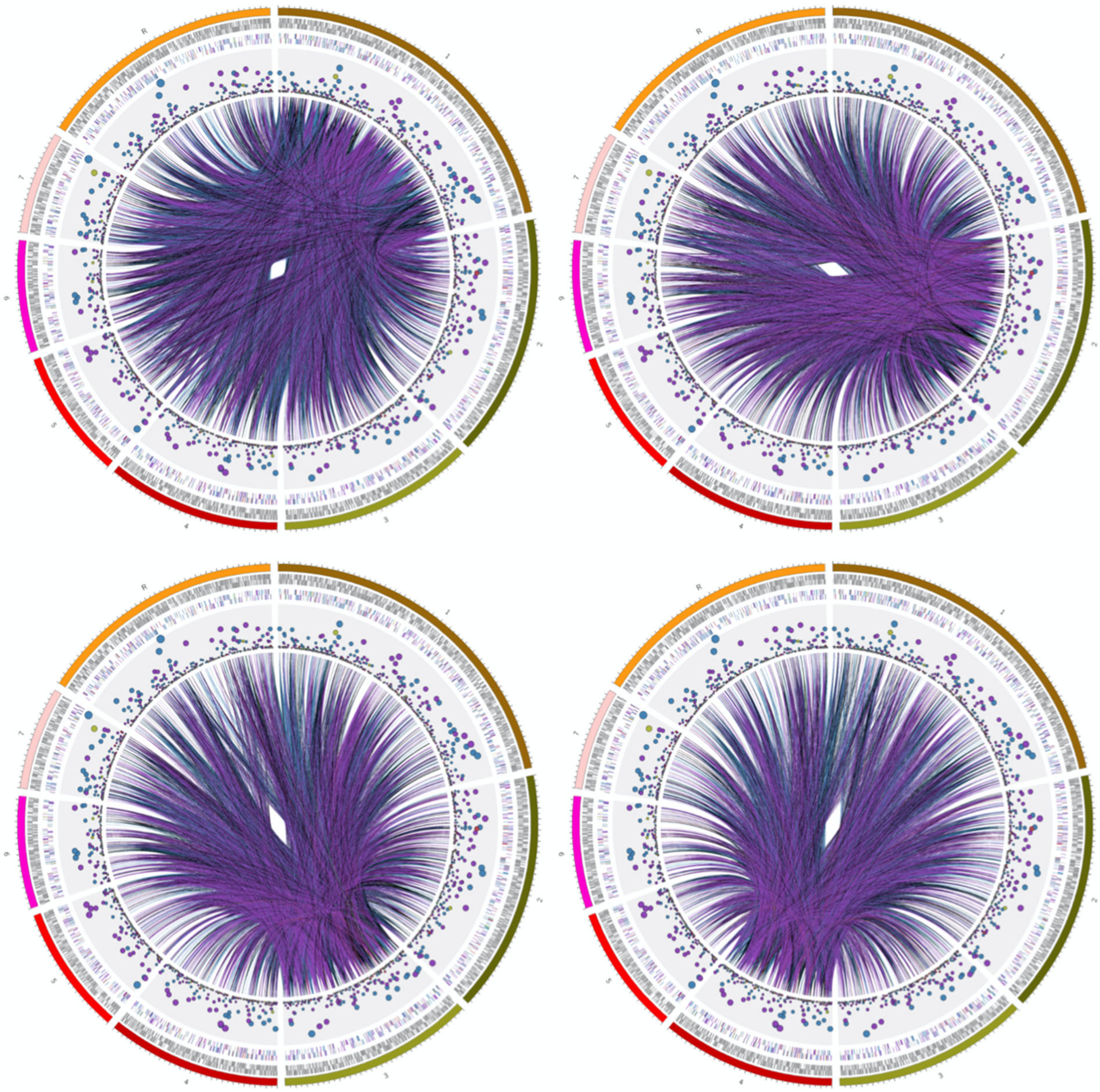

**Figure S21. Genes on chromosomes one through four are coevolving with genes on all other chromosomes in *Candida albicans*.** The first six tracks depict the same data as in Figure *N* (Genes on chromosomes one through four are coevolving with genes on all other chromosomes). Links depict genes that are coevolving with genes on a highlighted chromosome. Plots are arranged in ascending chromosome order from top left to bottom right.

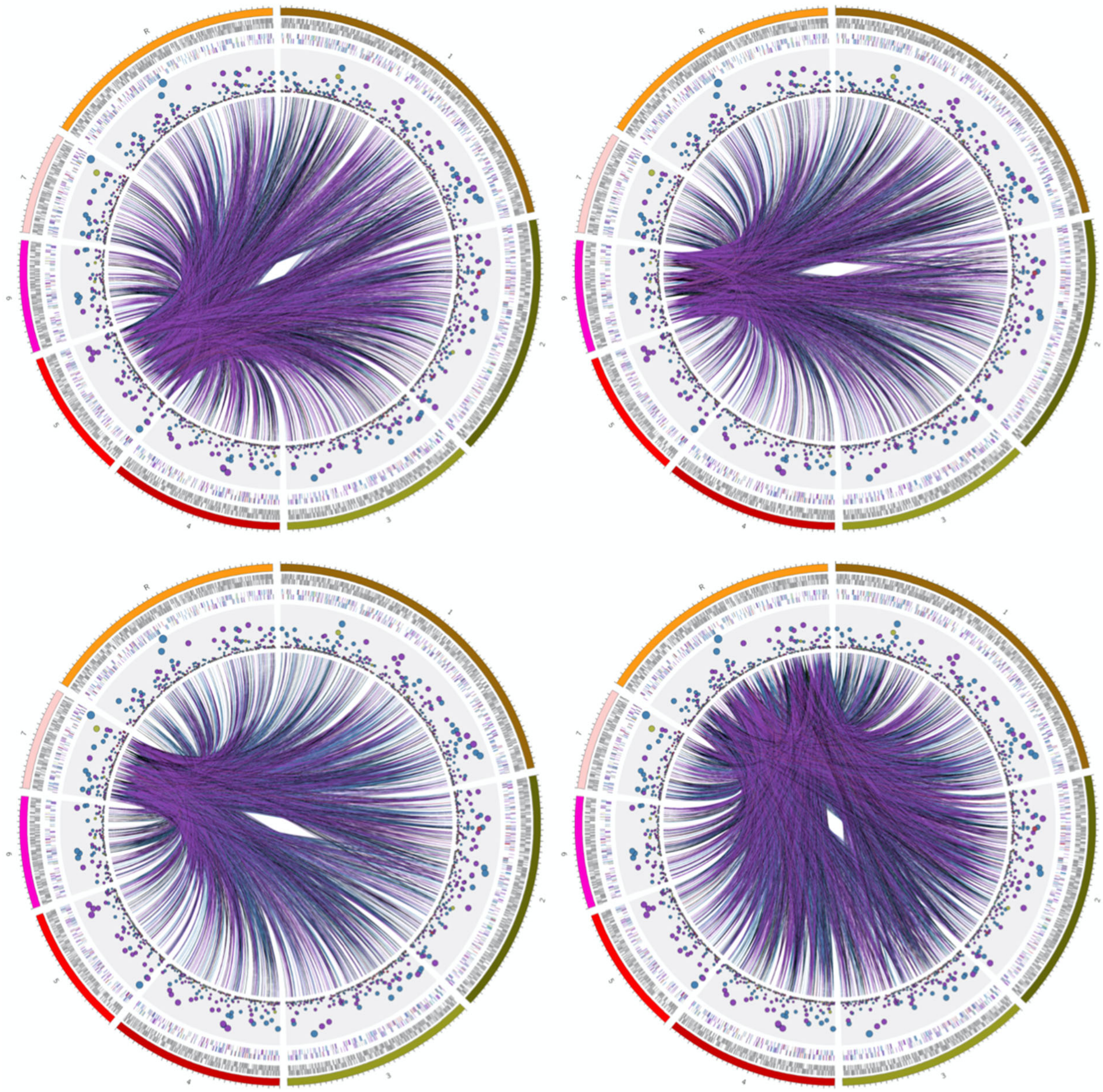

**Figure S22. Genes on chromosomes five through seven and chromosome R are coevolving with genes on all other chromosomes in *Candida albicans*.** The first six tracks depict the same data as in Figure *N* (Genes on chromosomes one through four are coevolving with genes on all other chromosomes). Links depict genes that are coevolving with genes on a highlighted chromosome. Plots are arranged in ascending chromosome order from top left to bottom right.

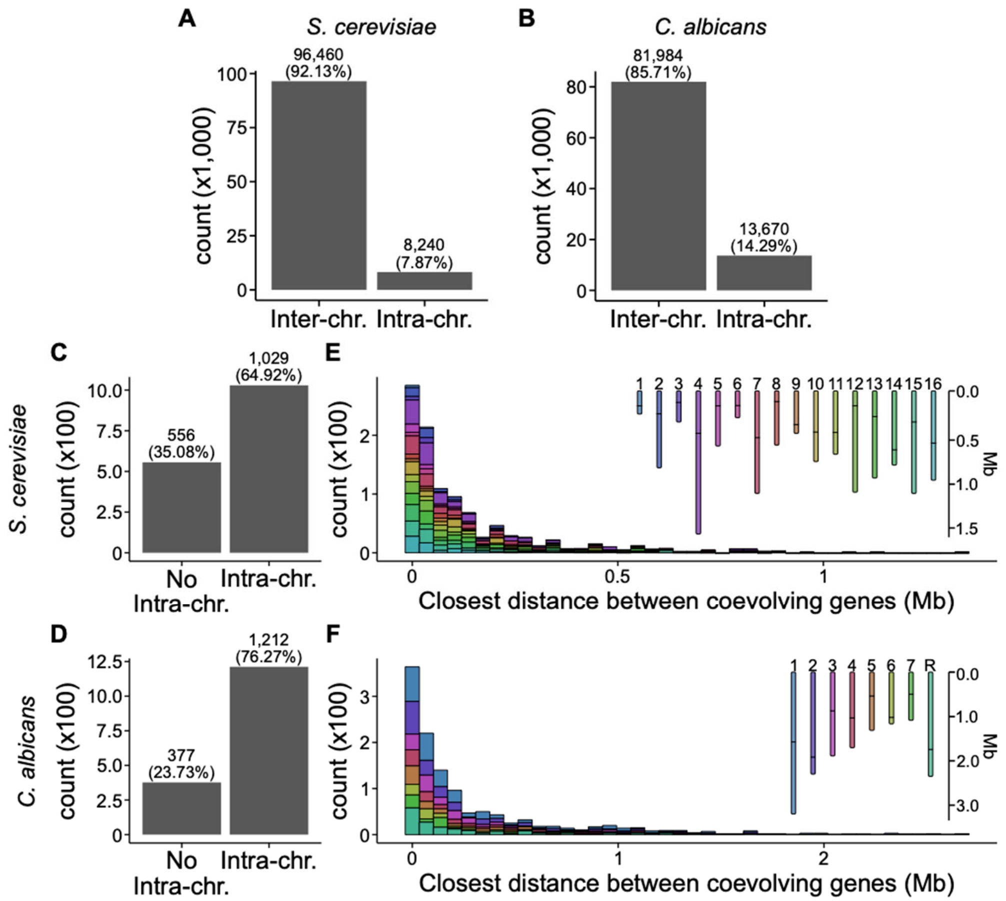


**Figure S23. There are many inter-chromosomal associations and intra-chromosomal associations can be long range.** (A and B) The total number of inter- and intra-chromosomal gene coevolutionary signatures reveal substantially more inter-chromosomal coevolution in comparison to intra-chromosomal coevolution in *S. cerevisiae* and *C. albicans*. (C and D) Examination of the number of genes that have no signatures of intra-chromosomal evolution and the number of genes with signatures of intra-chromosomal coevolution reveal a substantial portion of genes are not coevolving with genes on the same chromosome in either species of yeast. (E and F) The distribution of every gene and their closest coevolving partner among genes with evidence of intra-chromosomal coevolution reveals some genes may be coevolving despite substantial distances between them.

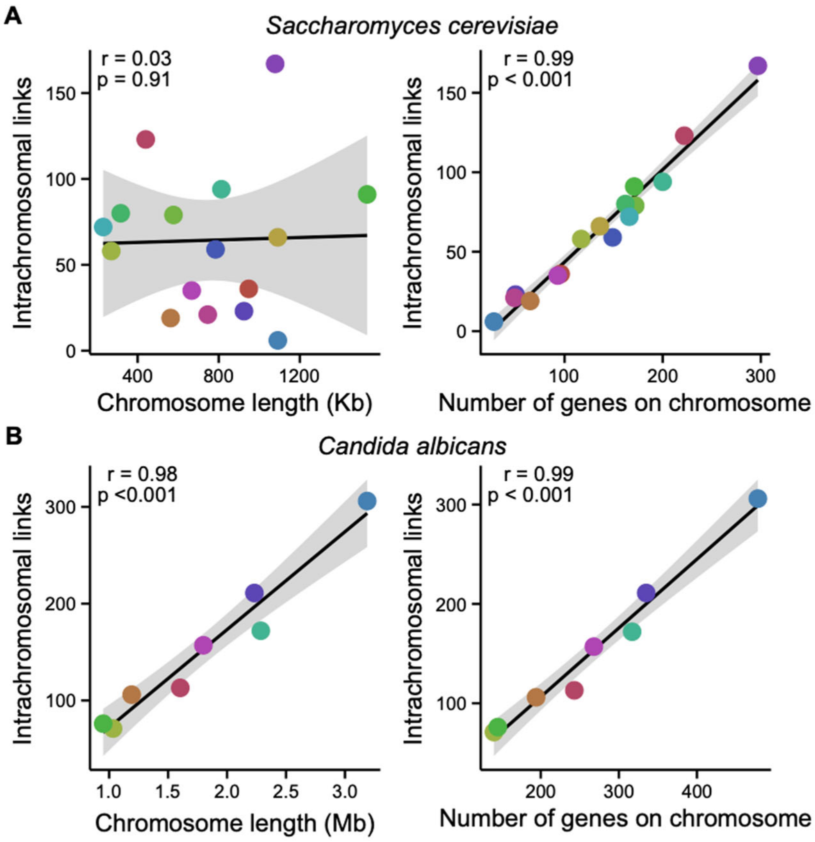


**Figure S24. The number of genes on a chromosome is the primary driver of the number of intrachromosomal links.** (A and B left panels) Chromosome length (x-axis) and the number of signatures of intrachromosomal gene coevolution (y-axis) are not significantly associated in (A) *S. cerevisiae* but are in (B) *C. albicans* ($r = 0.03$, $p = 0.91$ and $r=0.98$, $p<0.001$, respectively; Pearson Correlation). (A and B right panels) The number of genes on a chromosome (x-axis) and the number of signatures of intrachromosomal gene-gene coevolution (y-axis) are significantly associated for both (A) *S. cerevisiae* and (B) *C. albicans* ($r = 0.99$, $p < 0.001$ for both species; Pearson Correlation). Colors of each dot correspond to one chromosome. The color scheme is the same as depicted in Figure S21.

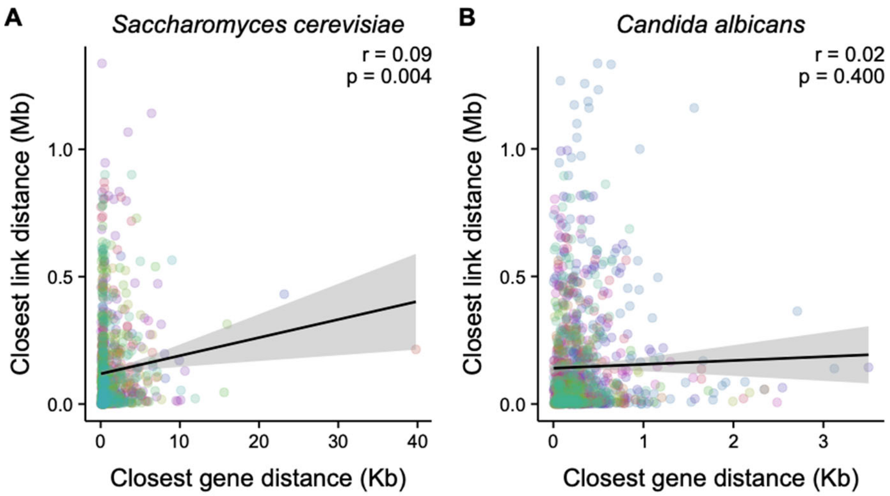


**Figure S25. The genomic distribution of orthologous genes is not driving the signature of long distance intra-chromosomal coevolution.** (A) In the genome of *S. cerevisiae*, there is no substantial association between a gene and the gene that it is most closely coevolving with (y-axis) and the closest gene in the data set (x-axis) ($r = 0.09$, $p=0.004$; Pearson Correlation). (B) Similarly, in *C. albicans*, there is not a significant association between the closest distance of a gene and an intra-chromosomal coevolving gene (y-axis) and the distance between the closest gene in the dataset (x-axis). Each data point represents a gene and the distance between either the closest intra-chromosomal gene that it is coevolving with (y-axis) and the distance between it and the closest gene in the data set (x-axis). Colors of each dot correspond to one chromosome. The color scheme is the same as depicted in Figure S21.

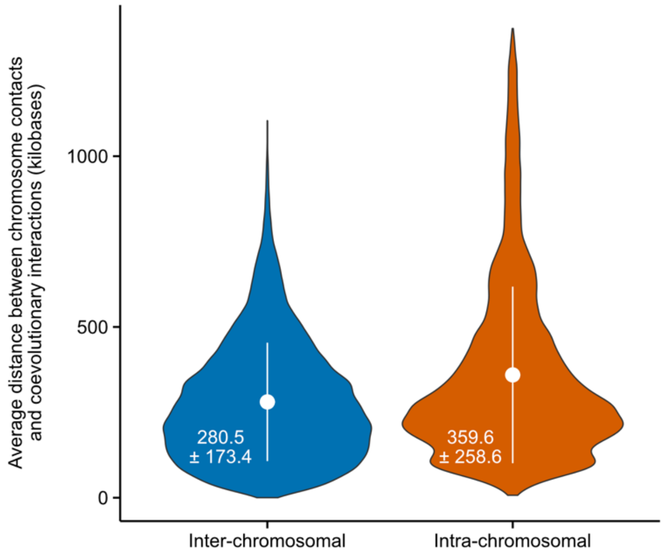


**Figure S26. Chromosome interactions do not seem to influence signatures of coevolution.** For every chromosomal interaction in *Saccharomyces cerevisiae*, the closest gene pair with a coevolutionary signature was identified. The average distance of the closest chromosomal interaction and coevolutionary signature was calculated and that distribution is shown here for interchromosomal (left) and intrachromosomal (right) interactions. White circles depict the mean and error bars are plus or minus one standard deviation. Mean and standard deviation values are shown in white text. Although some distances between chromosomal interactions and coevolving genes can be small, the average values observed are so great that chromosomal interactions do not seem to largely influence signatures of gene-gene coevolution.
